# Phosphoprotein Phosphatase Activity Positively Regulates Oligomeric Pyrin to Trigger Inflammasome Assembly in Phagocytes

**DOI:** 10.1101/2022.03.23.485108

**Authors:** Haleema S. Malik, Flora Magnotti, Nicole A. Loeven, Jose M. Delgado, Arminja N. Kettenbach, Thomas Henry, James B. Bliska

**Author notes:** Correspondence: James B. Bliska, Department of Microbiology and Immunology, Geisel School of Medicine at Dartmouth, 66 College Street, Hanover NH, 03755 USA.

## Abstract

Pyrin is a pattern-recognition receptor in phagocytes that triggers capase-1 inflammasome assembly in response to bacterial toxins and effectors that inactivate RhoA. Pyrin contains oligomerization domains and is negatively regulated by phosphorylation of two residues, S205 and S241 (murine) or S208 and S242 (human), via the kinases PKN1/2, which are activated by RhoA. Familial Mediterranean Fever (FMF) is caused by phagocyte production of pyrin gain of function variants, which have a lower threshold for inflammasome assembly upon RhoA-PKN axis inhibition. Inactivation of the RhoA-PKN axis removes negative regulation but a phosphoprotein phosphatase (PPP) is needed to positively regulate pyrin. No PPP that dephosphorylates pyrin has been identified, oligomerization of murine pyrin has not been studied, and the phosphorylation status of oligomeric pyrin is unknown. We used murine macrophages and FMF patient’s monocytes combined with the use of bacterial agonists and chemical inhibitors, native PAGE, phospho-specific antibodies and siRNA knockdowns to determine if a PPP positively regulates oligomeric pyrin. Results with broadly-specific inhibitors indicate that PPP activity is required to dephosphorylate murine and human pyrin in wild type or FMF patient’s phagocytes. Findings from native PAGE show that murine pyrin forms oligomers that are phosphorylated on S205 prior to RhoA inactivation. Inhibitors cause reduced mobility of murine pyrin on native PAGE and hyperphosphorylation of S242 in human pyrin, suggesting a PPP constitutively counterbalances PKN to keep the second site hypophosphorylated. Data from siRNA knockdown experiments implicate PP2A in dephosphorylation of S205 and positive regulation of pyrin in response to RhoA inactivation.

**Key points:** Murine pyrin is oligomeric and phosphorylated on S205 prior to inflammasome assembly

PPP activity positively regulates pyrin inflammasome assembly in mice and humans

The alpha and beta subunits of PP2A dephosphorylate murine pyrin S205 in macrophages

## Introduction

Numerous Gram-negative bacterial pathogens encode type III secretion systems (T3SSs) that are essential for virulence (1). T3SSs translocate effectors into or across the plasma membrane and into the eukaryotic cytosol during bacterial-host cell contact. Translocated effectors can inhibit innate immune responses to promote pathogenesis (1, 2). Host cells can sense perturbations caused by T3SSs and/or effectors and produce compensatory innate immune responses to counteract infection by pathogens (3, 4). Perturbations induced by T3SSs and/or effectors in macrophages infected with bacterial pathogens can result in the assembly of caspase-1 inflammasomes (5, 6). Inflammasomes are multimeric complexes that are assembled in response to various danger signals in the cell cytosol and serve as a molecular platform for the recruitment and maturation of caspase-1 into its active form (5, 6). Active caspase-1 cleaves gasdermin-D (GSDMD), pro-IL-1β and pro-IL-18 (7-9). The N-terminal domain of GSDMD forms pores in the plasma membrane which allow for release of mature IL-1β and IL-18 (7-9). GSDMD pores can also promote cell death, termed pyroptosis, resulting in the release of additional pro-inflammatory molecules (10). Pyroptosis amplifies the immune response by restricting growth of bacteria and facilitating bacterial clearance by neutrophils (7). Virulent bacterial pathogens can inhibit caspase-1 inflammasomes in host cells to counteract protective immune responses mediated by IL-1β, IL-18 and pyroptosis (11).

*Yersinia* species are Gram-negative bacterial pathogens that cause invasive human infections ranging in severity from plague (*Y. pestis*) to mesenteric lymphadenitis (*Y. pseudotuberculosis, Yptb*). These bacteria replicate in lymphoid tissues as extracellular microcolonies and require a plasmid-encoded T3SS for virulence (12). The T3SS delivers seven effectors called *Yersinia* outer proteins (Yops) into the cytosol of phagocytes that are in contact with *Yersinia* (12). The effectors inhibit key host cell processes, including phagocytosis and proinflammatory cytokine production (13). For example, YopE and YopT inhibit phagocytosis by inactivating Rho GTPases, including RhoA (13). YopE mimics a Rho GTPase activating protein (GAP) to accelerate GTP hydrolysis to GDP in RhoA (14), while YopT is a cysteine protease which cleaves RhoA at its C-terminus so that it is released from the membrane (15). Two effectors that are essential virulence factors inhibit inflammasome activation: YopM, a leucine-rich repeat (LRR)-containing protein (16-18) and YopK (19, 20). YopM inhibits the pyrin inflammasome (17, 18). YopK interacts with *Yersinia* translocon components to limit activation of NLRP3 and NLRC4 inflammasomes (19).

Inactivation of RhoA by YopE or YopT in macrophages triggers assembly of the pyrin inflammasome as a compensatory protective innate immune response (17, 18, 21, 22). Pyrin is a cytosolic pattern recognition receptor (PRR) which specifically senses pathogen induced modifications to RhoA (or RhoB or RhoC) to assemble an inflammasome (23). RhoA, RhoB and RhoC appear to be functionally redundant in regulating pyrin activity (23) and will be referred to as RhoA from here on. Inactivation of RhoA by the TcdB toxin of *Clostridioides difficile* or the TecA effector of *Burkholderia cenocepacia* also triggers pyrin inflammasome assembly in macrophages (23, 24). Pyrin is encoded by the *MEFV* gene (*Mefv* in mice) whose expression is restricted to phagocytes such as neutrophils, monocytes and activated macrophages (25). Pyrin is a TRIM protein and like other members of this family (26) human pyrin forms dimers via coiled coil (CC) domains (27). Additionally, human pyrin has been shown to form trimers (28). Pyrin potentially forms higher order oligomers (i.e. dimers of dimers), via B-box domains (21). It is not known if murine pyrin forms oligomers. Structurally, murine and human pyrin are similar except for their C-terminus. Murine pyrin lacks the B30.2 domain present in human pyrin (29). However, the mechanism of pyrin regulation appears to be conserved in both species (30).

Negative regulation of pyrin occurs in a manner similar to the guard hypothesis observed in plants, whereby a plant disease resistance PRR recognizes modifications to host proteins made by pathogens and induces an innate response (31). More specifically, pyrin follows an indirect guard hypothesis in which it does not directly interact with but is negatively regulated by RhoA (23). Hence, pyrin acts as the guard of RhoA (guardee) to protect host cells from bacterial effectors or toxins which target it. RhoA functions through a switch like mechanism between an “on” (Rho-GTP) and “off” (Rho-GDP) state. RhoA has numerous roles in phagocytosis, cell cycle and migration and is targeted by multiple bacterial pathogens (23, 32). Pyrin is negatively regulated by serine phosphorylation, via the kinase PKN (also known as PRK), which is in turn is activated by RhoA. PKN phosphorylates murine pyrin on serines 205 and 241 (S208 and S242 in human pyrin) within 14-3-3-binding sites located in the linker region to negatively regulate pyrin (30, 33). Mutational analysis of S205 and S241 by alanine substitution in ectopically expressed murine pyrin indicates that loss of either site results in partial inflammasome activation and when both sites are disrupted there is an additive effect (30). Similar studies of S208 and S242 in human pyrin show that the inflammasome is activated when either site is substituted (33, 34). Moreover, an S242R mutation in *MEFV* causes an autoinflammatory disease, PAAND, which leads to a partially active form of pyrin and further underscores the importance of linker phosphorylation in pyrin regulation (35). When pyrin is phosphorylated, the N-terminal pyrin domain (PYD) may be sequestered, preventing interaction with ASC (21). RhoA inactivation removes the negative regulation axis of pyrin, presumably resulting in dephosphorylation of S205/S208 and S241/S242 (21, 22, 30, 33). These findings lead to a model wherein when both serines in the pyrin linker are dephosphorylated, 14-3-3 detaches, the PYD is exposed, and ASC and caspase-1 are recruited to assemble inflammasomes (21, 30, 33). A limitation of this model is that no phosphoprotein phosphatase (PPP) that dephosphorylates pyrin has been identified and the phosphorylation status of pyrin oligomers is unknown.

Assembly of the pyrin inflammasome in phagocytes infected with WT *Yersinia* is blocked due to the action of YopM (17, 18). YopM binds to and activates the protein PKN and a second kinase, RSK (36). YopM hijacks PKN and RSK to keep pyrin phosphorylated (17, 21, 34). A *Yptb* Δ*yopM* mutant is fully virulent in Mefv^−/-^ mice, demonstrating that YopM promotes virulence by inhibiting pyrin (17). The importance of pyrin in pathogenesis extends beyond *Yersinia. C. difficile* and *B. cenocepacia* as well as numerous other bacteria including *Clostridium botulinum, Vibrio parahaemolyticus* and *Histophilus somni* secrete toxins that trigger assembly of this inflammasome in phagocytes (23).

Codon changes in the *MEFV* gene (e.g. M694V) that result in gain of function pyrin variants are responsible for the monogenic human autoinflammatory disease Familial Mediterranean Fever (FMF) (25, 34). FMF is characterized by recurrent episodes of fever and serositis and primarily affects eastern Mediterranean populations (37). Magnotti et al demonstrated that pyrin dephosphorylation was sufficient to promote inflammasome activation using PKN inhibitors on monocytes from FMF patients, which implies that these patients have a lower threshold for pyrin activation than healthy patients (38). It has been hypothesized that the high carrier frequency of FMF in Mediterranean and Middle Eastern populations is the result of a selective advantage in resistance to an unknown infection (34). Population genetic evidence is consistent with the idea that plague selected for FMF mutations in Middle Eastern populations (34). In addition, infection assays with human cells and mice provided evidence that pyrin FMF variants promote resistance to *Y. pestis* (34). These findings underscore the importance of the pyrin inflammasome for human health. Therefore, it is important to understand the dual role of pyrin in promoting immune responses against pathogens and autoinflammation in FMF and a deeper understanding of pyrin inflammasome regulation will aid in this goal.

Here we used primary murine macrophages, human THP-1 cells and primary monocytes from FMF patients in conjunction with bacterial toxins or effectors or chemical inhibitors, native PAGE, immunoblotting with phospho-specific pyrin antibodies and siRNA knockdowns to determine if PPP activity is required to dephosphorylate oligomeric pyrin and positively regulate inflammasome assembly in response to inactivation of the RhoA-PKN axis.

## Materials and Methods

### Bacterial strains

*Y. pseudotuberculosis* (*Yptb*) 32777 (serogroup O1) strains used in this study are as follows: wild-type (39), Δ*yopM* (40), Δ*yopK* (41), *yopE*^*R144A*^Δ*yopM* and *yopT*^*C139A*^Δ*yopM* (17). *Yptb* was grown at 28°C on Luria broth (LB) agar plates or with shaking in LB broth, supplemented with antibiotics when necessary. The *B. cenocepacia* strains used were AU1054 (WT) and Δ*tecA* mutant (42). *B. cenocepacia* was grown at 37°C on LB agar plates or with shaking in LB broth.

### Bone marrow isolation and cell culture conditions

Bone marrow derived macrophages (BMDMs) were obtained from femurs of 8-week-old WT C57BL/6 mice (Jackson Laboratories) and/or *Mefv*^*-/-*^ mice (43) as previously described (19). Cells were grown at 37°C in a humidified incubator in macrophage growth media (MGM) made up of Dulbecco’s modified Eagle Medium (DMEM) plus GlutaMAX (Gibco) containing 10% heat inactivated fetal bovine serum (FBS) (GE Health Care), 10% L929 cell-conditioned medium, 1 mM sodium pyruvate (Gibco), 10 mM HEPES (Gibco) for a week. On day 7 differentiated macrophages were divided into six-well plates at a density of 0.8×10^6^ cells/well in a total volume of 3 ml in MGM diluted to contain 10% L929 cell-conditioned medium (MGM 10/10) and primed for ∼18hrs with 100 ng/ml O26:B6 *Escherichia coli* LPS (Sigma).

### BMDM infection, intoxication, and inhibitor treatments

For BMDM infections, *Yptb* strains were grown overnight in LB at 28°C. The following day, cultures were diluted 1:40 in LB containing 20 mM sodium oxalate and 20 mM MgCl2 and grown at 28°C for 1 hr, then shifted to 37°C for 2 hr. Cultures were centrifuged and the bacterial pellets were resuspended in phosphate-buffered saline (PBS). Bacterial suspensions were then diluted in media lacking FBS and containing 10% L929 cell-conditioned medium (MGM 0/10). LPS-primed BMDMs were washed once with PBS and left uninfected or infected in MGM 0/10 with *Yptb* strains cultured under conditions as described above at an MOI of 30. Plates were centrifuged for 5 min at 106xg to facilitate contact between *Yptb* and macrophages. Plates were then incubated at 37°C in 5% CO_2_. In experiments where the glucosyltransferase toxin TcdB from *Clostridium difficile* (List Biological Laboratories, Inc.) was used as a positive control for pyrin inflammasome activation, BMDMs were intoxicated with TcdB at a concentration of 0.1LJμg/ml for 90min in MGM 0/10 media.

In experiments where phosphatase inhibitors were used and BMDMs were pre-treated for 15min with either 1.4mM DMSO as vehicle control or calyculin A (10nM diluted from 100μM stock, LC Laboratories), cyclosporin A (1μM diluted from 1mM stock, Sigma), okadaic acid (100nM or 1μM diluted from 1mM stock, Tocris), or tautomycetin (1μM or 2μM diluted from 1mM stock, Tocris) in MGM 0/10. MGM 10/10 was used for BMDMs pre-treated for 3hr with 100nM okadaic acid. For infection following pre-treatment, MGM 0/10 media containing bacteria and DMSO or the respective phosphatase inhibitors was added to the BMDMs, followed by centrifugation and incubation as above. Post-infection, cell supernatants were collected and processed for IL-1β detection by ELISA and cell lysates were prepared and processed for immunoblot analysis (see below).

### THP-1 cells, infection, and inhibitor treatment

THP-1 cells (ATCC TIB-202) were grown in filter sterilized RPMI 1640 media (Cat. No 30-2001) containing 10% heat inactivated fetal bovine serum and 0.05 mM 2-mercaptoethanol in 25cm^2^ flasks. When confluent the THP-1 cells were expanded in 75cm^2^ flasks. No cells were used that were passaged more than 30 times. For infections, THP-1 cells were seeded at a density of 0.8×10^6^ cells per well in 1.5ml of media in 6-well plates and pretreated with 1.4mM DMSO vehicle control or 10nM calyculin A for 15 min. Overnight (16-hour) cultures of *B. cenocepacia* were subcultured 1:100 in fresh LB on infection day and shaken at 37°C until the cultures reached mid-log phase (OD_600_ = 0.300). Cultures were then centrifuged, the LB was removed, and the bacterial pellets were resuspended in PBS and then diluted to an MOI of 20 in warmed RPMI 1640 media without FBS and containing DMSO or calyculin A. 1.5ml of supplemented media was added to each well and mixed by pipetting gently up and down. Following incubation for 3 hours at 37°C, the contents of each well was collected and centrifuged for 10 minutes at 14,000rpm and 4°C. The cell pellet was collected and lysed for immunoblot analysis as described below.

### FMF monocytes and inhibitor treatment

Primary monocytes purified as described (38) from three FMF patients carrying the p.M694V/p.M694V mutation were seeded into 96-well plates at 5×10^3^ cells/well, in RPMI 1640, GlutaMAX medium (Thermo Fisher Scientific) supplemented with 10% fetal calf serum (Lonza). Cells were incubated for 2.5h in the presence of LPS (10 ng/ml, InvivoGen) and, when indicated, pre-treated for 30 min with calyculin A (40nM, Sigma 208851), followed by treatment with UCN-01 (12.5 μM, Sigma) for 90 minutes. Following the incubation, cells were centrifuged, and supernatants were collected.

### U937 cell line and genetic manipulation

The protocol for genetically manipulating the human myeloid cell line U937 (CelluloNet, Lyon, France) was followed as described previously in (38). Cells were grown in RPMI 1640 medium with glutaMAX-I supplemented with 10% (vol/vol) FCS, 2 mM L-glutamine, 100 IU/ml penicillin, and 100 μg/ml streptomycin (Thermo Fisher Scientific). *MEFV*^KO^ cell lines generated by CRISPR/Cas9-mediated gene invalidations have been previously described (38, 44). p.M694V *MEFV* was cloned into the GFP-expressing plasmid pINDUCER21 (45) under the control of a doxycycline-inducible promoter through the pENTR1A (Invitrogen) vector using a synthetic DNA fragment (GeneArt) encoding the p.M694V Pyrin protein. Lentiviral particles were produced in 293T cells using pMD2.G and psPAX2 (from Didier Trono, Addgene plasmids #12259 and #12260), and pINDUCER-21 p.M694V Pyrin. U937 *MEFV*^KO^ cell lines were transduced by spinoculation and sorted at day 7 post-transduction based on GFP expression on an Aria cell sorter. Pyrin expression was induced by treatment with doxycycline (1 μg/ml) for 16 h before stimulation.

### Real-time cell death assay

U937 *MEFV*^KO^ cells expressing the p.M694V *MEFV* variant were seeded at 5×10^4^ per wells before stimulation in black 96-well plate (Costar, Corning) with propidium iodide (PI, Sigma) at 5 μg/ml. Three technical replicates per conditions were done. Cells were pre-treated with calyculin A (Sigma, 208851) at 40 nM for 30 min and then treated with UCN-01 (Sigma, U6508) at 12.5 μM. Real-time PI incorporation was measured every 5 min for 6 hours post-UCN-01 addition on a fluorimeter (Tecan) using the following wavelengths: excitation 535 nm (bandwidth 15 nm) and emission 635 nm (bandwidth 15 nm) (46, 47). Cell death was normalized (previously described, (38)) using PI incorporation in monocytes treated with 1% Triton X-100 for 15 min (=100% cell death). As a further correction, the first time point of the kinetics was set to 0.

### Protein analysis by SDS-PAGE and immunoblotting

To obtain cell lysates, BMDMs and THP-1 cells were lysed in mammalian protein extraction reagent (MPER, Thermo Scientific) along with cOmplete Mini (Roche) protease inhibitor and PhosSTOP (Roche) phosphatase inhibitor. The lysis buffer for THP-1 cells was supplemented with 10 nM calyculin A. Protein concentration was normalized after measurement with a bichinonic acid assay (BCA, Thermo Fisher Scientific) and 3-10μg of total protein cell lysates was run on 4 to 12% NuPAGE Bis-Tris SDS-PAGE gels (Invitrogen by ThermoFisher Scientific) and transferred to polyvinylidene difluoride membranes (ThermoFisher Scientific) using an iBlot 2 gel transfer device (Life Technologies). Membranes were blocked in 5% nonfat dairy milk (add company) and incubated with primary antibodies overnight. Primary antibodies used in this paper were: rabbit-anti-mouse monoclonal total pyrin antibody (1:1000 dilution, ab195975; abcam), rabbit-anti-mouse pyrin polyclonal (1:1000 dilution, (43)), rabbit-anti-mouse phospho-serine 205 monoclonal antibody (1:1000 dilution, ab201784; abcam), rabbit-anti-mouse phospho-serine 241 monoclonal antibody (1:1000 dilution, ab201784; abcam), rabbit-anti-mouse/human IL-1β (1:1000 dilution, number 12242; Cell Signaling), rabbit-anti-mouse/human polyclonal β-actin (1:1000 dilution, number 4967; Cell Signaling), mouse-anti-mouse/human/rat monoclonal PP1a (1:1000 dilution, number MA5-17239, ThermoFisher Scientific), rabbit-anti-*Y*.*pestis* polyclonal YopM (1:1000 dilution, provided by Susan Straley), mouse-anti-mouse monoclonal PKN (1:1000 dilution, number 393344, Santa Cruz Biotechnology), rabbit-anti-mouse monoclonal PP2A (1:1000 dilution, #2038, Cell Signaling), and rabbit-anti-mouse GSDMD (1:1000 dilution, ab209845, Abcam). Horseradish peroxidase-conjugated anti-rabbit or mouse antibody (Jackson Laboratory) were used as secondary antibodies. Protein bands bound by antibodies were visualized using chemiluminescent detection reagent (GE Healthcare) on an iBright FL1500 (ThermoFisher Scientific). Immunoblot quantification was performed using the iBright analysis software and ‘local corrected volume’ in the software was used to represent signal intensity for a band where indicated.

### Protein analysis by blue native (BN)-PAGE and immunoblotting

BN-PAGE immunoblotting was performed as described by Kofoed and Vance (48). All reagents for this procedure were obtained from Invitrogen except as stated. BMDMs were lysed in non-denaturing 1X Native-PAGE Novex sample buffer with the indicated concentrations of either digitonin (5% stock) or DDM (10% stock), MPER (Thermo Scientific) or RIPA lysis or extraction buffer (Thermo Scientific), along with cOmplete Mini (Roche) protease inhibitor and PhosSTOP (Roche) phosphatase inhibitor. Lysates were centrifuged at 20,817xg for 10min to obtain soluble and insoluble fractions. In some experiments the insoluble fractions of 1% digitonin lysates were analyzed by SDS-PASGE and immunoblotting as described above. Protein concentrations of soluble fractions were normalized after measurement with a bichinonic acid assay (BCA, Thermo Fisher Scientific) and 10μg or more of total protein cell lysates was mixed with NativePAGE G-250 additive and loaded on 4-16% NativePAGE Novex 3–12 % Bis-Tris gels. NativeMark unstained protein standard was used for molecular weight standard visualization. XCell SureLock Mini-Cell was used for running the gels. Samples were run at room temperature (RT) at constant 150V for 105-120min using pre-chilled buffers. Dark blue cathode buffer (10ml NativePAGE running buffer, 10ml NativePAGE cathode additive, 180ml deinonized water) was used for the inner chamber 12ntil samples had run up to 1/3^rd^ of the gel length after which the buffer was replaced with light blue buffer (10ml NativePAGE running buffer, 1ml NativePAGE cathode additive, 189ml deinonized water) for the remainder of the run. The outer chamber was filled with NativePAGE running buffer. Gels were soaked in 2X NuPAGE transfer buffer (Invitrogen) for 15min at shaking at room temperature. Proteins were transferred to polyvinylidene difluoride membranes (ThermoFisher Scientific) using the template program P3 (20V constant, 7min) and an iBlot 2 gel transfer device (Life Technologies). Post transfer membranes were incubated in 8% acetic acid for 15min with shaking, followed by ponceau S staining for 5min. Membranes were washed with distilled water for 5min to remove excess ponceau S stain then air-dried, reactivated by immersion in methanol, followed by blocking in 5% non-fat dry milk. Primary and secondary antibodies used are mentioned above.

### Protein analysis by two-dimensional blue native (BN)-PAGE and immunoblotting

The procedures used followed a previously described protocol (48). Samples were prepared as described above and separated on NativePAGE Novex 3–12 % Bis-Tris gels. Gel slices containing samples of interest were excised, trimmed to be 3mm shorter in length than the size of the 2D well in NuPAGE 4–12 % Bis-Tris 2D Gels, then submerged and shaken at RT in 1X NuPAGE sample buffer containing 0.1M DTT. Gel slices wwere then microwaved for ∼20sec to denature proteins followed by cooling with shaking at 10min at RT. Slices were then fitted into the 2D well and SDS-PAGE immunoblotting was performed as described above.

### siRNA knock-down

siRNAs targeting *Pkn1* (s115927), *PP2Aca* (#1: s72067, #2: s72066) and *PP2Acb* (s72069) were obtained from Invitrogen. For siRNA gene knock-down experiments, 400pmol of *Pkn1*,1000pmol of *PP2Aca* siRNAs, and for combined PP2Ac knock down, 500pmol each of PP2Aca siRNA and PP2Acb siRNA were transfected into BMDMs (1×10^6^ cells) by electroporation using the Neon Transfection System (Invitrogen) and electrical parameters of 1400V, 20ms and 2 pulses. BMDMs were then replated in 6-well plates and grown in MGM 10/10 media. 24hr following transfection, media was changed to fresh MGM 10/10 and LPS was added for overnight incubation. 48hr following transfection BMDMs were infected with Δ*yopM Yptb* as described above.

### Quantitative real-time PCR (RT-qPCR)

At 48hr following transfection, uninfected BMDMs were lysed with trizol reagent (Invitrogen) to isolate RNA according to the manufacturer’s instructions. 500-1000ng of RNA was used to make cDNA using SuperScript IV First-Strand Synthesis System (Invitrogen). Approximately ∼5ng of cDNA was quantified using PowerUp SYBR Green Master Mix (Invitrogen) in the (add machine name). Primer specificity was checked by melt-curve analysis. Expression of target genes was normalized to the expression of the housekeeping gene Hprt. For fold change analysis data was transformed using the ‘relative standard curve method and comparative threshold cycle (Ct) method (ΔΔCt)’ as described by Applied Biosystems. Published primer sequences were used for qPCR analysis (49) and the sequences are: Hprt, forward 5′-TGAAGT ACTCATTATAGTCAAGGGCA-3′ and reverse 5′-CTG GTGAAAAGGACCTCTCG-3′; Ppp2ca, forward 5′-TCTTCCTCTCACTGCCTTGGT-3′ and reverse 5′-CAG CAAGTCACACATTGGACCC-3′; Ppp2cb, forward 5′-AAGGCGTTCACCAAGGAGCT-3′ and reverse 5′-ACAGCGGACCTCTTGCACAT-3′.

### IL-1β quantification

IL-1β in BMDM supernatants was quantified using a murine (MLB00C; R&D Systems) and human (R&D Systems) ELISA kit by following the manufacturer’s instructions.

### Statistical analysis

GraphPad Prism was used to perform statistical analyses. IL-1β ELISA data were analyzed by one-way or two-way analysis of variance (ANOVA; Bonferroni multiple-comparison test), or by Krukal-Wallis test (Dunn’s multiple comparison test) as specified.

### Ethics statement

The study was approved by the French Comité de Protection des Personnes (CPP,#L16-189) and by the French Comité Consultatif sur le Traitement de l’Information en matière de Recherche dans le domaine de la Santé (CCTIRS, #16.864). The authors observed a strict accordance to the Helsinki Declaration guidelines. All FMF patients fulfilled the Tel Hashomer criteria for FMF and were homzygous for the p.M694V mutation.

## Results

### PPP activity is required for pyrin dephosphorylation and positive regulation of inflammasome assembly

Inactivation of RhoA removes negative regulation by PKN, however, a PPP is needed to positively regulate pyrin by serine dephosphorylation. There are seven members in the PPP family: PP1, PP2A, PP2B (also known as calcineurin or PP3), PP4, PP5, PP6, and PP7 (50). LPS-primed bone marrow-derived macrophages (BMDMs) were pretreated for 15 min with DMSO as a vehicle control, or a PPP inhibitor to test the role of PPPs in dephosphorylation of pyrin. The following PPP inhibitors were used: calyculin A (CA) at 10 nM and okadaic acid (OA) at 1000 nM inhibit PP1, PP2A, PP4, PP5, and PP6 but not PP2B; OA at 100 nM more specifically inhibits PP2A; and 1000nM of cyclosporin A (CSA) inhibits PP2B. After the pretreatment the BMDMs were maintained in the presence of DMSO or the inhibitors during intoxication for 90 min with *C. difficile* TcdB, which glucosylates and inactivates RhoA (51), to trigger pyrin inflammasome assembly (23). BMDM lysates were analyzed by immunoblotting with monoclonal antibodies specific for total pyrin or phosphoserine 205 (PS205) (30) or actin as loading control. We were unable to monitor PS241 in murine pyrin because the antibody reported to recognize this modification (30) did not work in our hands for this purpose but did recognize human pyrin PS242 (see Fig.1C). We found that treatment of BMDMs with CA or 1000 nM OA prevented pyrin S205 dephosphorylation in response to TcdB intoxication (Fig.1A). Results of ELISA on BMDM supernatants showed that CA or 1000 nM OA also significantly reduced release of mature IL-1β during TcdB intoxication (Fig.1B). CSA treatment did not inhibit pyrin S205 dephosphorylation or IL-1β release upon TcdB intoxication (Fig.1AB) which indicates that PP2B does not regulate the pyrin inflammasome. 100nM OA treatment was also ineffective (Fig.1AB) but previous publications (52) and subsequent experiments (see Fig.6 below) indicate that at this concentration longer pretreatments are needed for the inhibitor to reach effective concentrations in cells. We noted that CA and 1000nM OA in the absence or presence of TcdB caused an upshift in the positions of pyrin bands on the immunoblots, but not a corresponding increase in PS205 signal (Fig.1A). This suggests that S205 is uniformly phosphorylated and a PPP constitutively dephosphorylates pyrin at an additional site(s), possibly S241, and when this phosphatase is inhibited, hyperphosphorylation by PKN results in reduced mobility of pyrin by SDS-PAGE. In addition, these results suggest that a PPP other than PP2B is required to dephosphorylate pyrin S205 and trigger assembly of the inflammasome in response to TcdB.

**Fig.1.**
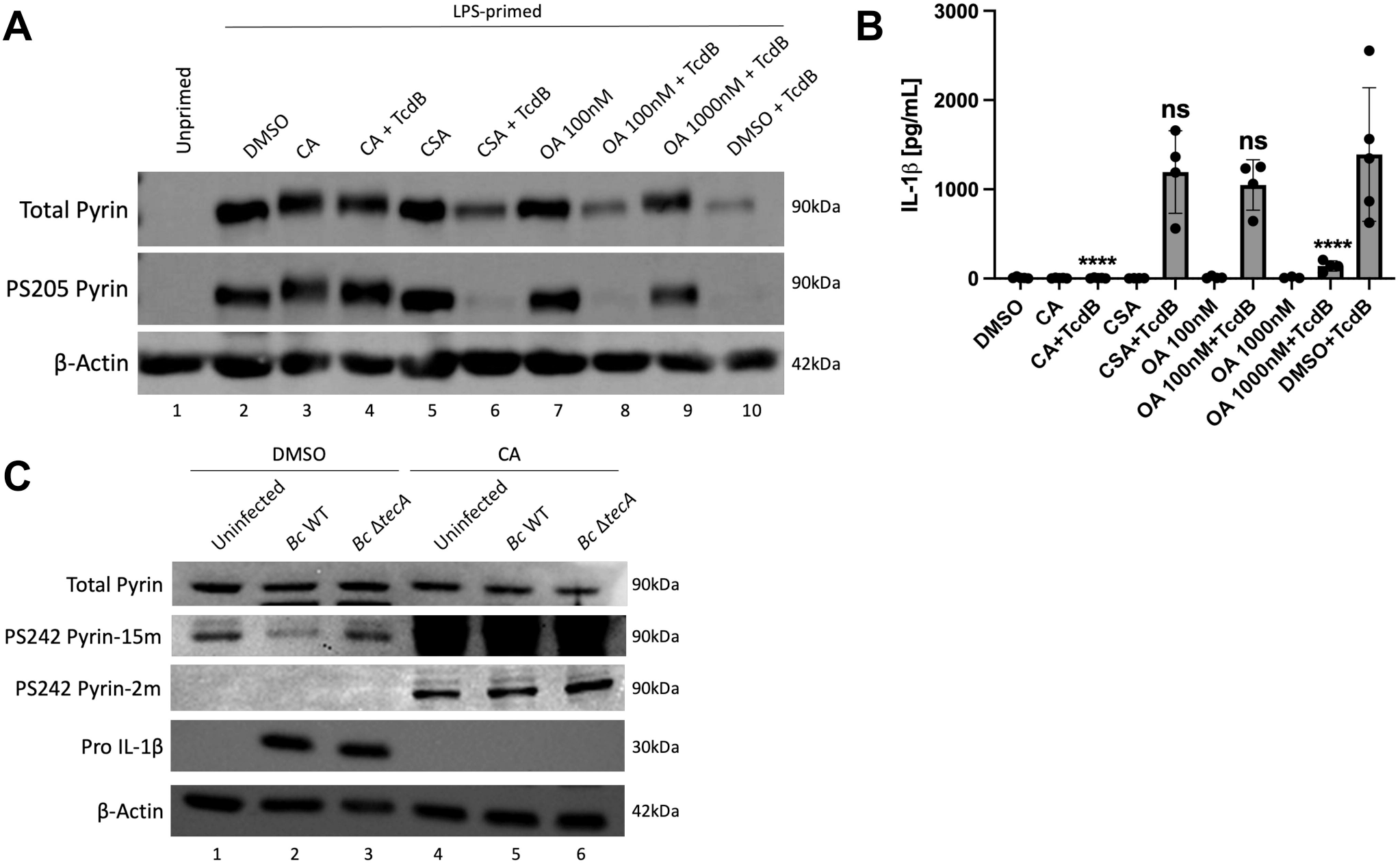
PPP inhibition with CA or OA prevents pyrin dephosphorylation and reduces IL-1β release. LPS-primed BMDMs were pretreated for 15mins and maintained with either 10nM calyculin A (CA), 100nM okadaic acid (OA), 1000nM OA or 1000nM cyclosporin A (CSA), and left unintoxicated or intoxicated with 0.1μg/mL of TcdB for 90 mins (A-B). Unprimed BMDMs or treatment with vehicle DMSO were used as controls. A) Immunoblot analysis of total and PS205 pyrin in BMDM lysates. Actin was used as a loading control. B) Mature IL-1β in supernatants as quantified by ELISA. In (B) Two-way ANOVA with Bonferroni post hoc correction was applied to calculate significance and p-values as compared to DMSO+TcdB are indicated. P-value<0.05 was considered significant; <0.0001 (****); not significant (ns). Each data group is presented as an average (error bars are standard deviation) of at least three independent experiments. Naïve THP-1 cells were pre-treated for 15 min and maintained with either DMSO or 10nM CA, and either left uninfected or infected with *B. cenocepacia* (Bc) WT or Δ*tecA* for 180 min at an MOI of 20 (C). Immunoblot analysis of total and PS242 pyrin and pro-IL-1β in THP-1 lysates is shown. PS242 pyrin signals are show with long (15m) or short (2m) exposures for comparison. Actin was immunoblotted as a loading control.

We used CA treatment to determine if a PPP is important for YopE and YopT to trigger pyrin S205 dephosphorylation and inflammasome assembly in BMDMs infected with *Yptb*. BMDMs treated with DMSO or CA as above were infected with a Δ*yopM* mutant to activate the pyrin inflammasome. We also infected DMSO or CA-treated BMDMs with a Δ*yopK* mutant, which triggers assembly of the NLRP3 inflammasome (and to a lesser degree the NLRC4 inflammasome) (19, 20), since there is evidence that PP2A is important for dephosphorylation and activation of NLRP3 (49). Immunoblot and ELISA analysis of BMDMs pre-treated with CA followed by Δ*yopM* infection showed that pyrin remained phosphorylated on S205 and there was a significant decrease in IL-1β release (Fig.2AB). Results of infection with Δ*yopK* also showed reduced IL-1β release when the BMDMs were treated with CA, although the decrease was not significant (Fig.2B). Pyrin was not dephosphorylated on S205 in response to Δ*yopK* infection of untreated BMDMs as expected (Fig.2A). The finding that CA had a stronger inhibitory effect on IL-1β secretion with Δ*yopM* as compared to Δ*yopK* infection (Fig.2B) might reflect residual NLRC4 inflammasome assembly triggered by the latter strain (19).

**Fig.2.**
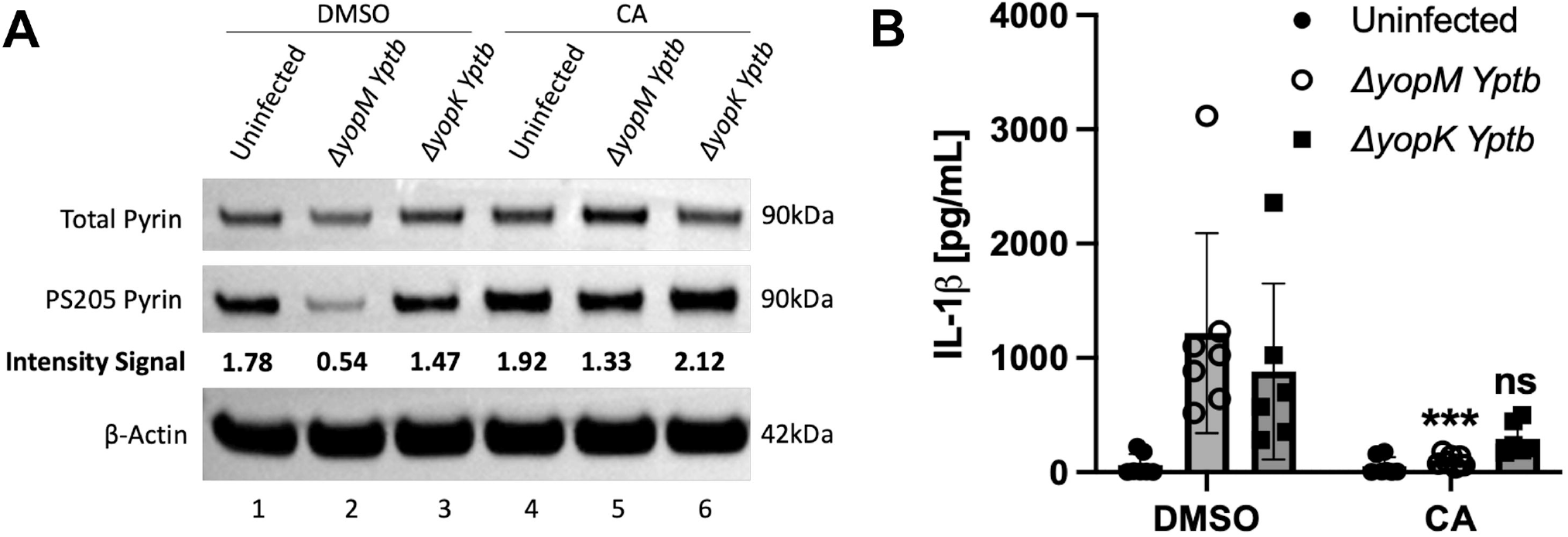
PPP inhibition with CA reduces pyrin dephosphorylation and IL-1β release in BMDMs infected with Δ*yopM Yptb*. LPS-primed BMDMs were pre-treated for 15mins and maintained with either DMSO or 10nM CA, and infected with the Δ*yopM* or Δ*yopK* strain at an MOI of 30 for 90 mins. Treated uninfected BMDMs were analyzed in parallel. A) Immunoblot analysis of PS205 and total pyrin in BMDM lysates. Actin was used as a loading control. Quantified immunoblot band intensity signals from one experiment representing PS205 pyrin/total pyrin are indicated below the respective blot image. B) Mature IL-1β in supernatants as quantified by ELISA. Two-way ANOVA with Bonferroni post hoc correction was applied to calculate significance and p-values of each treatment as compared to its corresponding DMSO treated infected strain is indicated. P-value<0.05 was considered significant; <0.001 (***); not significant (ns). Each data group is presented as an average (error bars are standard deviation) of at least three independent experiments.

Similar results were obtained when CA was used to treat BMDMs infected with Δ*yopM Yptb* strains expressing only active YopE (*yopT*^*C139A*^) or YopT (*yopE*^*R144A*^) (Fig.S1AB). In addition, immunoblotting indicated that CA did not decrease steady state levels of pro-IL-1β (Fig.S1A).

*B. cenocepacia* infection of naive THP-1 cells has been shown to trigger assembly of the pyrin inflammasome (53). To determine if PPP inhibition prevents dephosphorylation of human pyrin we infected THP-1 cells with *B. cenocepacia* WT or Δ*tecA* strains in the absence or presence of DMSO or CA. Lysates were immunoblotted with antibodies that recognize total pyrin, PS242, pro-IL-1β or actin as loading control. *B. cenocepacia* infection triggered dephosphorylation of S242 in a TecA-dependent manner in DMSO-treated THP-1 cells (Fig.1C). Interestingly, when THP-1 cells were pretreated and maintained in the presence of CA, S242 became hyperphosphorylated in both uninfected and infected conditions (Fig.1C), which was evident from comparing the short and long exposures of the immunoblots. The hyperphosphorylation of pyrin in the presence of CA in uninfected THP-1 cells suggests that a PPP is constitutively counterbalancing PKN to keep S242 hypophosphorylated. CA treatment prevented infection-induced production of pro-IL-1β as determined by immunoblotting (Fig.1C) and as a result the impact of the inhibitor on release of mature IL-1β from the THP-1 cells could not be measured. These data indicate that a PPP dephosphorylates pyrin S242 in response to inactivation of RhoA by TecA in THP-1 cells, but it appears that a PPP is acting constitutively to keep most of the pyrin in the cell dephosphorylated at this site prior to infection.

PKN inhibitors activate pyrin inflammasome assembly and pyroptosis in monocytes from FMF patients with the M694V mutation (38). The effect of CA on release of IL-1β from FMF patient’s monocytes activated by treatment with the PKN inhibitor, UCN-01, was measured. Pre-treating LPS-primed FMF monocytes with 40 nM CA followed by UCN-01 treatment significantly inhibited IL-1β release as compared to UCN-01 treatment alone (Fig.3). Moreover, CA treatment also prevented UCN-01 mediated cell death as measured by propidium iodide influx in a human monocytic cell line, U937, expressing the p.M694V *MEFV* variant (Fig.S2). Thus, in an FMF context where the threshold of activation has been shown to be lower and relies on pyrin dephosphorylation (38, 54), a PPP is constitutively countering the effect of the PKN kinases and inhibition of PPP activity prevents inflammasome activation. Altogether these data replicate the inhibitory effect of CA as seen on murine pyrin dephosphorylation and activation in BMDMs in human cells.

**Fig.3.**
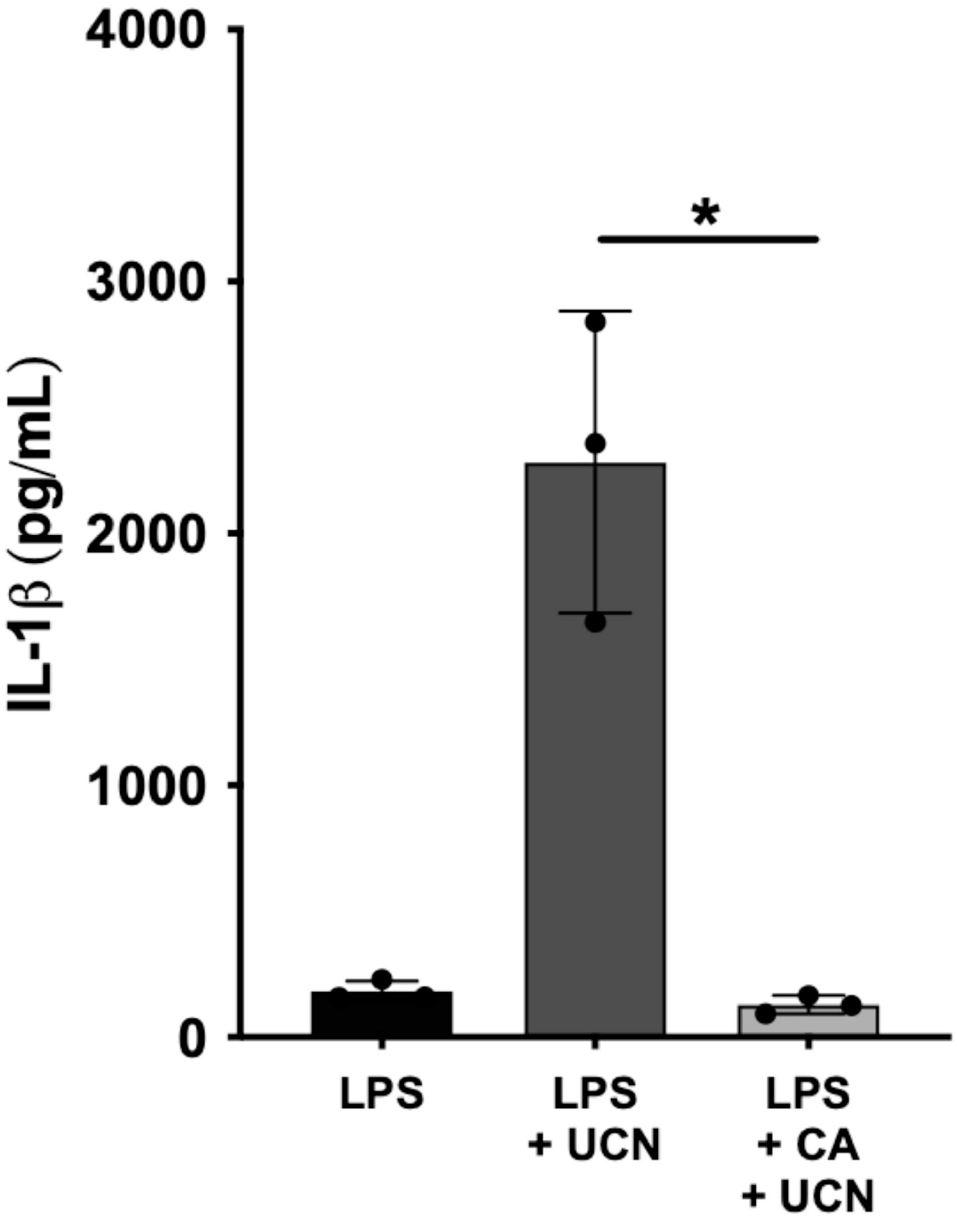
CA inhibits UCN-01-mediated IL-1β release from FMF patient’s monocytes. Monocytes from three FMF patients were primed first with LPS (10ng/ml) during 2.5h and, when indicated, pre-treated with CA (40 nM) during 30 min. Cells were then treated with 12.5 μM UCN-01 for 90min. Mature IL-1β in supernatants as quantified by ELISA is shown. Each symbol presents an average (error bars represent median interquartile range) of three biological replicates for one patient. Kruskal-Wallis test with Dunn’s correction for multiple comparisons was applied. P-value less than 0.05 was considered significant; <0.05 (*).

To further delineate which PPP activities positively regulate pyrin, we used tautomycetin (TTN), a more specific inhibitor of PP1 (55, 56). WT BMDMs were pretreated for 15 min and maintained in DMSO or 1 or 2μM of TTN during infection with Δ*yopM Yptb* (Fig. S1CD). We found that treatment of BMDMs with TTN did not prevent pyrin PS205 dephosphorylation or IL-1β release in response to infection with Δ*yopM Yptb* (Fig. S1CD). Longer pretreatment of BMDMs with 1μM TTN also did not decrease IL-1β secretion upon Δ*yopM* infection (unpublished observation). Our combined results from treatment of BMDMs with various phosphatase inhibitors suggest that a PPP other than PP1 and PP2B is required to activate pyrin. *PPP activity regulates phosphorylation of oligomeric pyrin*

Although human pyrin has been shown to form trimers (28) the oligomeric status of murine pyrin has not been investigated. In addition, it is not known if pyrin oligomers are differentially phosphorylated or dephosphorylated. We used blue native PAGE (BN-PAGE) and immunoblotting (48, 57) to examine the impact RhoA inactivation and PPP inhibition on phosphorylation of oligomeric pyrin. To optimize the procedure, we first tested different non-ionic detergent extraction conditions on untreated uninfected WT or *Mefv*^−/-^ BMDMs. As shown in Fig.S3, when extracts of untreated BMDMs were separated by BN-PAGE and immunoblotted for total (A) or PS205 (B) pyrin, we detected oligomers of pyrin in WT but not *Mefv*^*-/-*^ BMDMs. Monomeric pyrin (∼90 kDa) was not detected and based on the apparent molecular weight of the oligomers we tentatively assigned them as dimer (∼242 kDa) and higher-order oligomers (∼480-720 kDa). Thus, most of the pyrin in BMDMs is in a higher oligomeric form and phosphorylated on S205. Recovery of total and PS205 pyrin was optimal using 0.5% or 1% digitonin (Fig.S3) and 1% was used going forward.

We next analyzed the phosphorylation and oligomeric status of pyrin in BMDMS infected with WT or Δ*yopM Yptb* in the absence of CA using BN-PAGE and immunoblotting. As shown in Fig.4, the total (A) and PS205 (B) pyrin oligomer signals decreased in BMDMs infected with Δ*yopM* (lane 3). Loss of the total protein signal was largely due to increased insolubility of the dephosphorylated form of pyrin in non-ionic detergent (Fig.4E) as we have documented previously (22), although some degradation of pyrin upon Δ*yopM* infection is possible. This insolubility issue prevented us from determining if the oligomeric state of pyrin is altered upon dephosphorylation. In BMDMs treated with CA there was an upshift in the positions of the dimeric and higher oligomeric forms under all conditions, most evident with the PS205 signal, and there was no insolubility or dephosphorylation of pyrin in response to infection with Δ*yopM* (Fig.4CD). In addition, in the presence of CA a very high molecular weight oligomer at ∼1048 kDa was detected with the PS205 signal under all conditions (Fig.4D). These results confirm the upshift in mobility due to hyperphosphorylation of pyrin in the presence of CA seen in some SDS-PAGE immunoblots (Fig.1A). These data further suggest that hyperphosphorylation of pyrin increases oligomerization.

**Fig.4.**
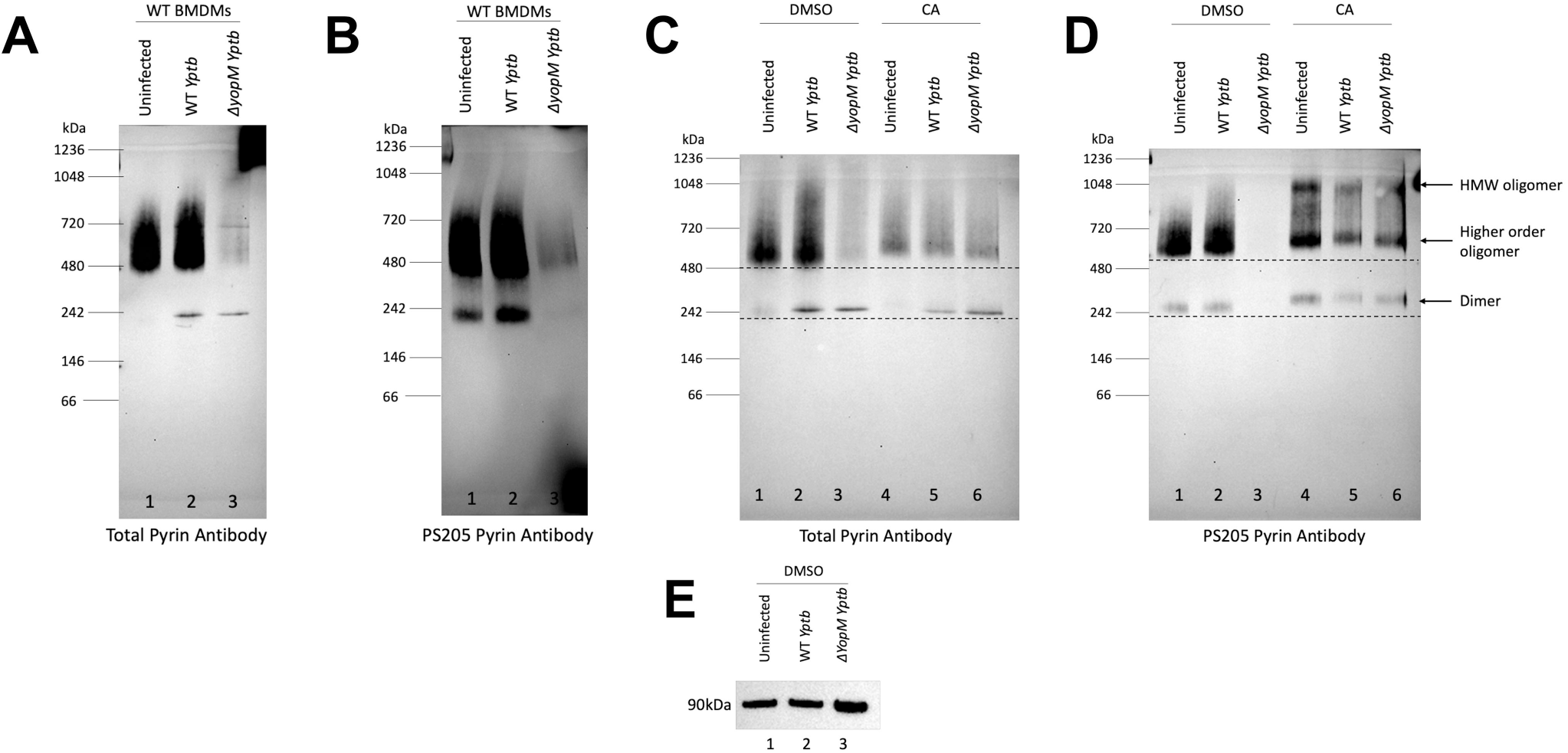
Impact of PPP inhibition with CA and *Yptb* infection on pyrin oligomers and phosphorylation as determined by BN-PAGE and immunoblotting. WT BMDMs left untreated (A,B) or pre-treated and maintained with DMSO or 10 nM CA (C-E) were left uninfected or infected with WT or Δ*yopM Yptb* at an MOI of 30 for 90 mins. BMDM lysate samples prepared in 1% digitonin were separated by BN-PAGE and immunoblotted for total (A,C) or PS205 (B,D) pyrin. BN-PAGE MW standards are shown on the left and tentative positions of pyrin oligomers are shown on the right. Dashed horizontal lines in (C,D) aid viewer appreciation of band upshifts in the presence of CA. Insoluble proteins from 1% digitonin lysates of DMSO treated samples were analyzed by SDS-PAGE and total pyrin immunoblotting (E).

### YopM interacts with oligomeric pyrin

YopM can interact with pyrin as determined by co-immunoprecipitation (17, 18, 34). We wondered if YopM interacts with the different oligomeric forms of YopM and if this interaction is sensitive to PPP inhibition. To determine if YopM is associated with pyrin oligomers, samples from untreated WT BMDMs left uninfected or infected with WT or Δ*yopM Yptb* were separated by BN-PAGE and immunoblotted with antibody to YopM. We detected a YopM signal at the dimer and higher-order oligomer positions in the WT but not in the Δ*yopM Yptb* samples (Fig.5A, lanes 2 and 3). We then repeated the experiments using WT BMDMs untreated or treated with CA or untreated *Mefv*^*-/-*^ BMDMs. In WT BMDMs there was a YopM signal corresponding to dimeric pyrin in the DMSO and CA treated WT *Yptb* infected samples (Fig.5B, lanes 2 and 5). The higher order oligomeric YopM signal around ∼480kDa seen with DMSO treatment was not present in the CA treated sample (Fig.5B, lane 5). In addition, there was a faint YopM signal around ∼480kDa in the *Mefv*^*-/-*^ BMDMs infected with WT *Yptb* (Fig.5B, lane 8). Two dimensional (2D) BN-PAGE immunoblot analysis of the DMSO treated and WT *Yptb* infected BMDMs sample showed a good correspondence in the molecular weights of the YopM and oligomeric pyrin signals (Fig.5C). These results strongly indicate that YopM interacts with dimeric and higher order oligomers of pyrin. Furthermore, when the CA treated and WT *Yptb* infected sample was subjected to 2D BN-PAGE analysis, the YopM signal around ∼480kDa disappeared but the band around ∼242kDa was still present (Fig.5D), recapitulating the BN-PAGE results shown in Fig.5B. Possible explanations for why YopM interaction with the higher order pyrin oligomer is sensitive to PPP inhibition, and the presence of a weak YopM signal at this position in *Mefv*^−/-^ BMDMs are discussed below (see Discussion).

**Fig.5.**
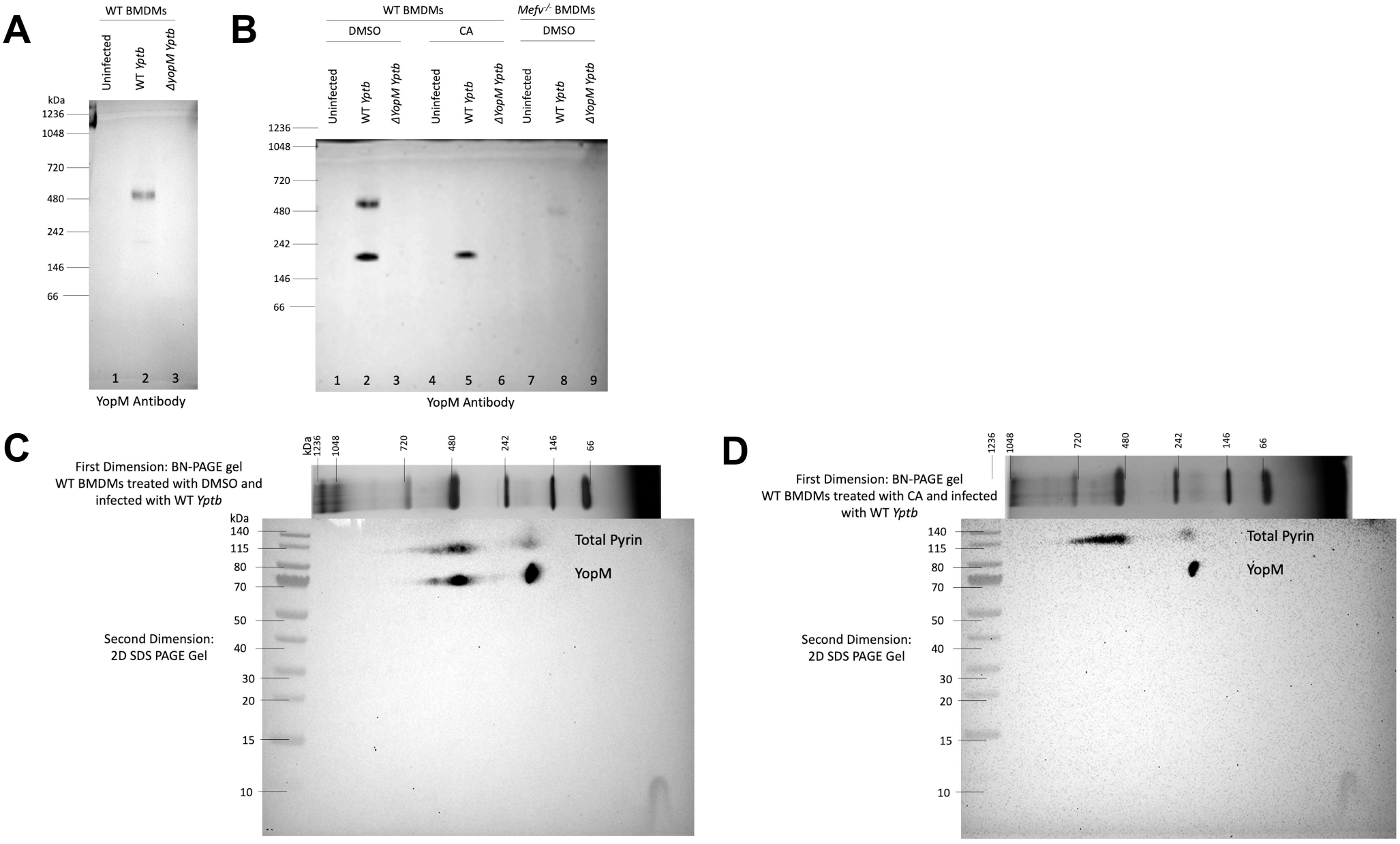
Detection of YopM interaction with pyrin oligomers by one-dimensional and two-dimensional BN-PAGE and immunoblotting. Untreated WT BMDMs (A) or WT or *Mefv*^*-/-*^ BMDMs pre-treated for 15 min and maintained in DMSO or 10nM CA (B-D) were either left uninfected or infected with WT or Δ*yopM Yptb* at an MOI of 30 for 90 mins. Cell lysates were solubilized in 1% digitonin, separated by one dimensional BN-PAGE and immunoblotted for YopM (A,B). BN-PAGE MW standards are shown on the left and tentative positions of YopM oligomers are shown on the right. 1% digitonin samples from WT BMDMS treated with DMSO (C) or 10nM CA (D) and infected with WT *Yptb* were separated in the first dimension by BN-PAGE after which the gel slice was isolated and analyzed by SDS-PAGE in the second dimension followed by immunoblotting for pyrin and YopM. MW markers for each dimension are indicated.

*Longer pretreatment with 100 nM OA reduces pyrin dephosphorylation and inflammasome assembly*

As mentioned previously, OA penetration inside cells is slow and using a lower concentration of OA generally requires longer pretreatment time to achieve sufficient inhibitory levels inside the cell. In contrast CA readily partitions inside cells (52, 58, 59). We found that WT BMDMs pretreated for 3hr and maintained with 100nM OA during infection with Δ*yopM Yptb* showed reduced pyrin dephosphorylation (Fig.6A) and a significant decrease in IL-1β release as compared to the DMSO control (Fig.6B); whereas 15min pretreatment of BMDMs with 100nM OA did not have such an inhibitory effect on pyrin activation (Fig.1AB). Given the selectivity of 100 nM OA to preferentially inhibit PP2A/PP4 over PP1/PP5 (59), these results suggest that PP2A or PP4 could be involved in positively regulating pyrin. Since most serine/threonine phosphatase activities inside the cell are carried out by PP1 and PP2A (50, 60) our results with TTN and longer pretreatment with OA in BMDMs allude to a role for PP2A in positive regulation of pyrin activation.

**Fig.6.**
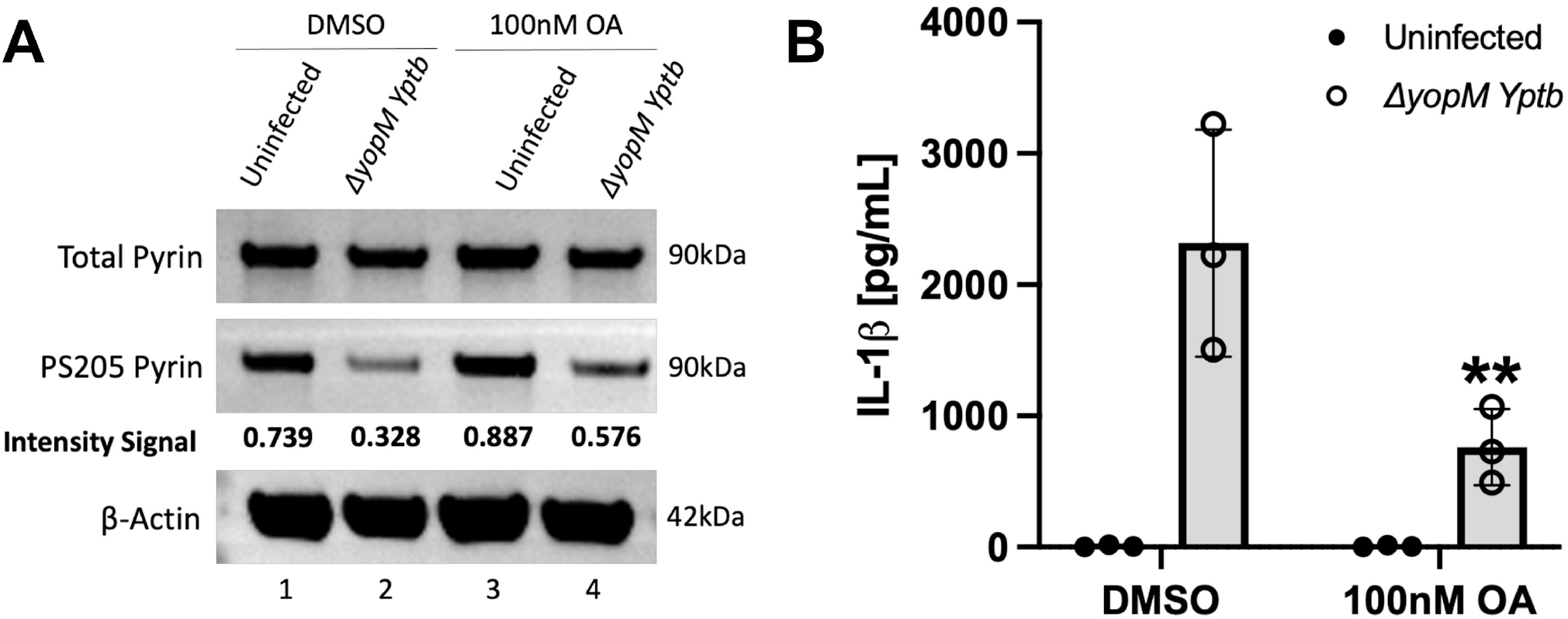
Longer pretreatment with 100nM OA reduces pyrin dephosphorylation and IL-1β release in BMDMs infected with Δ*yopM Yptb*. LPS-primed BMDMs were pre-treated for 3 hrs and maintained with either DMSO or 100nM OA and infected with the Δ*yopM Yptb* at an MOI of 30 for 90 mins. Treated uninfected BMDMs were analyzed in parallel. A) Immunoblot analysis of PS205 and total pyrin in BMDM lysates. Actin was used as a loading control. Quantified immunoblot band intensity signals from one experiment representing PS205 pyrin/total pyrin are indicated below the respective blot image. B) Mature IL-1β in supernatants as quantified by ELISA. Two-way ANOVA with Bonferroni post hoc correction was applied to calculate significance and p-value of OA treatment as compared to its corresponding DMSO treated and Δ*yopM* infected sample is indicated. P-value<0.05 was considered significant; <0.01 (**). Each data group is presented as an average (error bars are standard deviation) of at least 3 independent experiments.

### siRNA knockdown implicates PP2A in dephosphorylation of pyrin S205

We next tested the effect of knocking down PP2A in BMDMs on pyrin inflammasome assembly. PP2A has two catalytic subunit isoforms, alpha (PP2Aca) and beta (PP2Acb), which only differ in a few amino acids from each other. We first tested two different siRNAs targeting the mRNA of the alpha catalytic subunit of PP2A (49). The siRNAs (1000pmol) were electroporated into WT BMDMs which were also LPS-primed. PKN1 siRNA was used as a positive control (17, 33). RT-qPCR showed that there was significant and selective knock down of PP2Aca mRNA with both siRNAs as compared to the positive control and PP2Acb analyzed in parallel (Fig.S4A). The siRNA electroporations were then repeated and LPS-primed BMDMs were left uninfected or infected with Δ*yopM Yptb*. Immunoblotting showed that there was full depletion of PKN1 with the corresponding siRNA but only an apparent ∼50% knock down of PP2A with the siRNAs #1 and #2 (Fig.S4B). Since the PP2A antibody used detects both isoforms the remaining signal detected could have corresponded to PP2Acb. Upon Δ*yopM* infection there was no inhibition of dephosphorylation of pyrin at S205 with either PP2Aca siRNAs (Fig. S4B). However, PP2Aca #2 siRNA decreased IL-1β secretion (unpublished observation) which could indicate that this siRNA has an off-target effect when used at the high concentration. A PP2Acb siRNA when tested by itself at 500pmol selectively knocked down the corresponding mRNA transcripts and did not decrease IL-1β release as compared to the control (Fig.7A and Fig.S4C, respectively). We next tested the effect of a combined PP2A alpha and beta catalytic subunit knock down. BMDMs were electroporated with PKN1 or PP2Aca and PP2Acb siRNAs (500pmol each) and LPS primed as before. RT-qPCR showed that there was a decrease in the mRNA transcripts of both PP2A subunits (Fig.7A). Immunoblotting showed a greater depletion of PP2A protein as compared to knocking down the alpha subunit alone (compare Fig.S4B and Fig.7B). Upon Δ*yopM* infection PS205 pyrin was maintained and there was a significant decrease in IL-1β secretion with both siRNA combinations tested (Fig.7B and C, respectively). Together, these results implicate PP2A alpha and beta subunits as acting redundantly to dephosphorylate pyrin S205.

**Fig.7.**
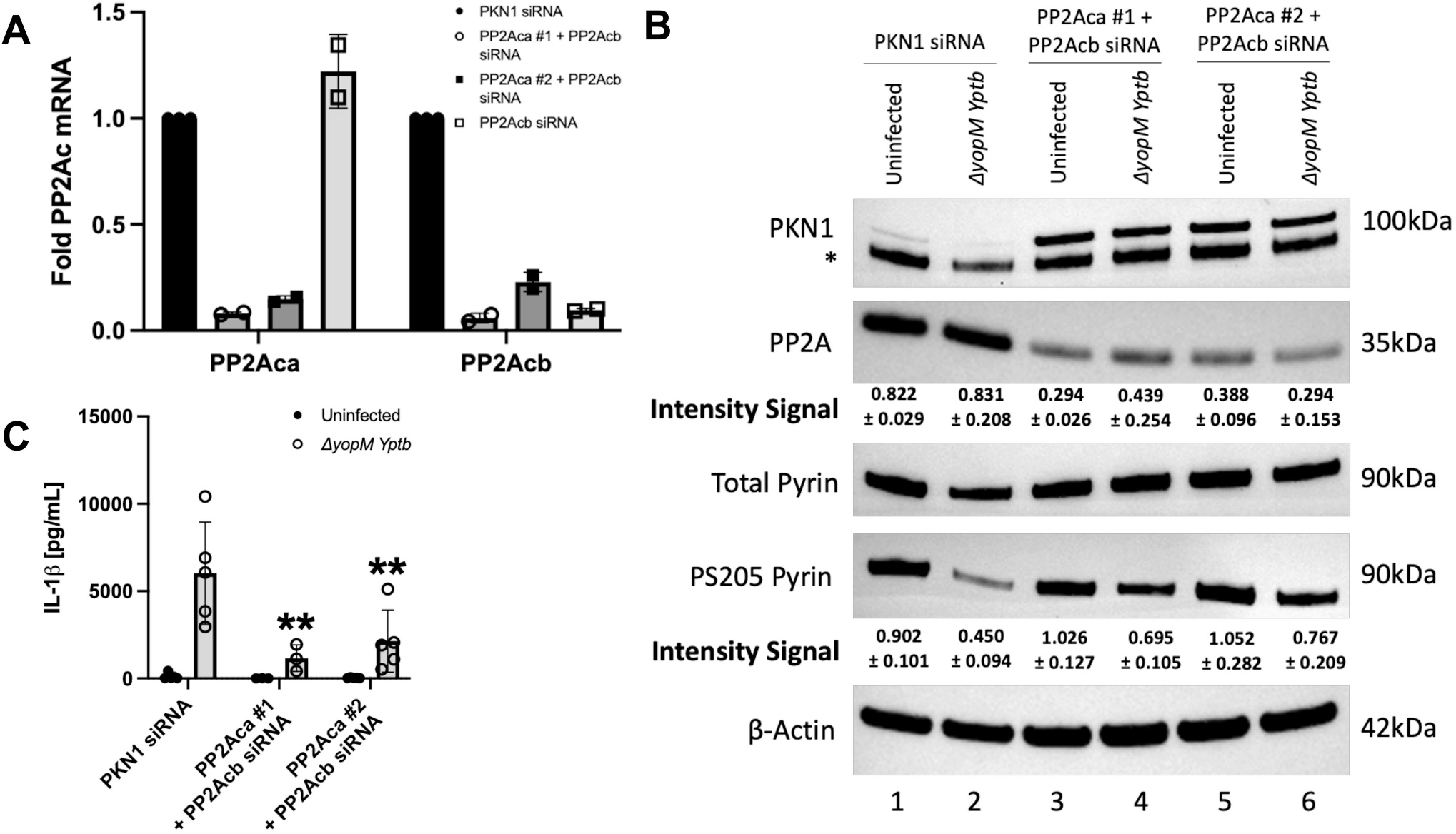
siRNA knock down of PP2A in BMDMs inhibits pyrin S205 dephosphorylation and IL-1β release. WT BMDMs were electroporated with siRNAs targeting PKN1 (400pmol) or PP2Aca + PP2Acb (500pmol each). 24hr after electroporation the BMDMs were LPS-primed. 48hr following electroporation, cells were either left uninfected or infected with Δ*yopM Yptb* at an MOI of 30 for 90 mins. A) RT-qPCR analysis of mRNA transcripts of PP2Aca and PP2Acb in uninfected BMDMs. Results were normalized to Hprt mRNA levels. Samples with error bars represent standard deviation for an average of two independent experiments. B) Immunoblot analysis of PKN1, PP2A, total and PS205 pyrin in uninfected and Δ*yopM Yptb-*infected BMDM lysates. Actin was used as a loading control. Asterisk (*) indicates bands for total pyrin on an immunoblot that was reprobed for PKN1. Quantified immunoblot band intensity signals from three independent experiments representing PP2A/β-Actin and PS205 pyrin/total pyrin are indicated below their respective blot images. Signals of bands from immunoblots from three independent experiments was averaged to calculate mean and standard deviation. C) Mature IL-1β in supernatants as quantified by ELISA. Two-way ANOVA with Bonferroni post hoc correction was applied to calculate significance and p-value of samples as compared to the PKN1 siRNA and Δ*yopM* infected sample is indicated. P-value<0.05 was considered significant; <0.01 (**). Each data group is presented as an average (error bars are standard deviation) of at least three independent experiments.

## Discussion

In this study, we aimed to understand if a PPP activity is important for positive regulation of inflammasome assembly by oligomeric pyrin. There are seven different PPPs in the mammalian cell: PP1, PP2A, PP2B (PP3), PP4, PP5, PP6 and PP7. In the human genome there are ∼428 Ser/Thr kinases but only ∼30 Ser/Thr phosphatases (60). The remarkable diversity of PPP substrates, substrate recognition and cellular localization is achieved by the association of catalytic subunits with non-catalytic subunits to form multimeric holoenzymes (50). The active sites of these phosphatases share 100% similarity, and most phosphatase inhibitors which bind to the PPP active sites are broadly specific for this reason.

CA is a relatively non-specific and hydrophobic PPP inhibitor. It can easily partition through cell membranes and potently inhibits most PPPs except for PP2B and PP7 (59). CA was found to inhibit both murine and human pyrin inflammasome activation. Using 10nM CA we obtained evidence that a PPP activity is important for pyrin S205 dephosphorylation and inflammasome assembly in BMDMs in response to RhoA inactivation by TcdB or YopE/T during Δ*yopM Yptb* infection. The same CA treatment also prevented TecA-dependent dephosphorylation of pyrin S242 in THP-1 cells infected with *B. cenocepacia*. It has been shown previously in mouse peritoneal macrophages that 50nM CA prevented activation of NLRP3, AIM2 and NLRC4 inflammasomes, indicating that PPP activity can be important for inflammasome activation in general, and is perhaps acting at a latter step (61). We observed that 10nM CA prevented infection-induced pro-IL-1β synthesis in THP-1 cells infected with *B. cenocepacia* illustrating that results with this inhibitor need to be carefully controlled and interpreted since it can block multiple steps in an inflammasome pathway. Moreover, 40 nM CA was also found to reverse the effect of a PKN inhibitor and potently inhibit pyrin activation in FMF patient monocytes. These cells have previously been demonstrated to exhibit a lower activation threshold for inflammasome activation as compared to monocytes from healthy donors, and pyrin dephosphorylation has been to shown to be sufficient to trigger inflammasome assembly in FMF monocytes (37, 38). It is possible that in the FMF disease context, a balance between the PKN kinases and a PPP regulates inflammasome assembly and inhibition of either shifts the equilibrium in favor of the other protein. Results from our experiments with treatment of BMDMs with CA and Δ*yopK Yptb* mutant infection showed that there was a non-significant decrease in IL-1β release which can be attributed to inhibition of NLRP3 but not of the NLRC4 inflammasome. Luheshi et al used a higher CA concentration of 50nM (61) which could explain why we didn’t observe inhibition of NLRC4 inflammasome in response to Δ*yopK* infection with 10nM CA treatment.

We were also able to show using BN-PAGE that murine pyrin forms oligomers consisting of dimers and higher order oligomers that are phosphorylated on S205 when pyrin is inactive. Due to the insolubility of active pyrin in non-ionic detergent we were unable to determine if dephosphorylation changes the oligomeric status of pyrin. However, PPP inhibition with CA resulted in the appearance of a very high molecular weight mystery oligomer which indicates that phosphorylation increases oligomerization of pyrin. Pyrin belongs to the TRIM family of proteins and Trim5α which is involved in HIV viral capsid recognition is known to form dimers and higher order lattices consisting of dimers of dimers or trimers of dimers (26). The higher order murine pyrin oligomers (∼480-720kDa) that we observe on BN-PAGE immunoblots could represent dimers of dimers or trimers of dimers similar to those formed by Trim5α. It has been previously shown through crosslinking studies in THP-1 cell lysates that human pyrin can form trimers which is mediated through its coiled coil domains (28). In the same study it was also shown that the B-box domain of human pyrin interacts with the N-terminal pyrin domain to sequester it and prevent interaction with ASC (28). However, Weinert et al reported that a truncated version of human pyrin forms an antiparallel dimer in vitro through interaction of coiled coil domains (27). These discrepancies in trimer vs dimer formation by human pyrin could be attributed to experimental and technical differences. It is expected that regulation of human and murine pyrin by oligomerization would be conserved but due to structural differences, it is possible that murine pyrin forms dimers and human pyrin forms trimers through their respective coiled coil domains. To rule out technical differences it would be informative to investigate oligomerization of human pyrin by BN-PAGE instead of crosslinking. The phosphorylation status of human pyrin oligomers should also be determined.

CA treatment resulted in an upshift in molecular weights of PS205 and total murine pyrin bands in all conditions on BN-PAGE immunoblots, as wells as on some SDS-PAGE immunoblots. In addition, CA treatment of THP-1 cells resulted in hyperphosphorylation of S242 in human pyrin. This indicates that a PPP acts constitutively on the second linker site in pyrin to dephosphorylate it and when this PPP is inhibited the second serine becomes hyperphosphorylated. These results suggest that one of the linker sites can be hypophosphorylated yet pyrin remains inactive. This finding is surprising given the data that phosphorylation of the second linker site is critical for negative regulation of pyrin, based on alanine substitution of this residue in ectopically expressed pyrin variants (33, 34). In addition, the S242R substitution causes PAAND in which the pyrin variant is considered constitutively active (35). Pull down experiments in this study also showed that 14-3-3 proteins are not bound to the S242R variant and that pyrin is dephosphorylated (35). Jeru et al. showed previously that a S208A pyrin variant has reduced 14-3-3 binding while the S242A mutant abolishes 14-3-3 binding (62). This suggested that 14-3-3 binding could follow a gatekeeper hypothesis in which PS242 is the dominant site which promotes 14-3-3 binding and PS208 is a secondary site (62). These findings may indicate that hypophosphorylation of the second linker site is sufficient to keep pyrin oligomers inactive. This could facilitate fast dephosphorylation of the hypophosphorylated second sites, resulting in rapid activation of pyrin in response to RhoA inactivation by pathogens. A caveat of this concept is our limitation of not being able to measure the phosphorylation status of S241 in murine pyrin under the conditions used.

We were also able to use one and 2D BN-PAGE and immunoblotting to detect interaction of murine pyrin with YopM during WT *Yptb* infection of BMDMs and found that this interaction was sensitive to PPP inhibition. The fact that YopM targets the oligomers argues that they are poised to become activated upon dephosphorylation. We hypothesize that the faint YopM signal on BN-PAGE immunoblots from the *Mefv*^*-/-*^ BMDMs infected with WT *Yptb* could result from self-oligomerization of the YopM protein. The YopM protein from *Y. pestis* is known to form tetramers in vitro (36, 63), so it is conceivable that the *Yptb* YopM protein can form higher order oligomers on its own in *Mefv*^*-/-*^ BMDMs. One explanation for the selective loss of the YopM signal at the position of the higher order oligomers from the CA-treated and WT *Yptb-*infected samples is that the affinity of YopM binding to pyrin is reduced by hyperphosphorylation.

OA in comparison to CA has a greater inhibitory constant to bind to the catalytic subunit of PP2A than of PP1 and is reported to enter cells slowly (59, 64). Our results indicated that 15min pretreatment with 1000nM OA is sufficient to inhibit S205 pyrin dephosphorylation and IL-1β release in BMDMs upon RhoA inactivation. Stutz et al reported that PP2A is important for NLRP3 activation through dephosphorylation using the same conditions and concentration of OA (49). We only observed a reduction in S205 dephosphorylation and a significant decrease in IL-1β secretion in response to Δ*yopM* infection when BMDMs were pretreated with 100nM OA for 3hr but not for 15min, which can be explained by the slow entry of the inhibitor. We hypothesize from these results that PP2A plays a role in dephosphorylation of S205 and positive regulation of the pyrin inflammasome in BMDMs.

PP2A, along with PP1, carry out most serine/threonine dephosphorylation events in cells and partake in many essential activities for the cell (50). PP2A is abundantly present in cells and forms heterotrimers consisting of a catalytic subunit, a scaffolding A subunit, and a regulatory B subunit (50). There are two isoforms of the catalytic and scaffolding subunits, and four isoforms of the regulatory subunits, which account for the diverse activities and substrates regulated by PP2A (60). Since it is an essential protein for the cell and very likely has a long half-life which explains why we were unable to fully deplete it in the cell. The alpha and beta catalytic subunit of PP2A only differ by a few amino acids and PP2Aca is more abundantly present in cells than PP2Acb. Our results indicated that siRNAs targeting PP2Aca did not reduce S205 pyrin dephosphorylation in BMDMs infected with Δ*yopM*. However, a combined knock down of both catalytic subunits of PP2A showed that S205 pyrin phosphorylation was maintained and there was a significant decrease in IL-1β secretion upon Δ*yopM* infection. These results indicate that the two catalytic subunits of PP2A act redundantly to dephosphorylate the pyrin linker. This is distinct from the case of NLRP3 where knockdown of the alpha catalytic subunit of PP2A alone was sufficient to prevent assembly of the inflammasome (49). To confirm our results, additional siRNAs targeting the alpha and beta catalytic subunit of PP2A should be tested. In addition, the impact of PP2A knockdowns on PS241 in murine pyrin and PS208/242 in human pyrin needs to be investigated. The phosphorylation status of murine PS241 and human PS208 in pyrin could be investigated using immunoprecipitation combined with mass spectrometry or the use of pan-specific antibodies to phosphoserine. Additionally, the use of pyrin variants with amino acid changes that ablate or mimic serine phosphorylation (30) could be helpful to corroborate the siRNA knockdown data. Moreover, in a broader disease context, the effect of PP2A knockdown on FMF pyrin activation should also be explored because it may offer insights for new therapeutic strategies.

PPPs recognize and bind to regulatory proteins or to substrates via short linear motifs (SLiMs) (50). For example, the B56 regulatory subunit of PP2A binds through LxxIxE motifs to its substrates (50). Interestingly, murine pyrin contains two LxxIxE motifs in the coiled-coil domain and human pyrin has one LxxIxE motif in the linker. Since both murine and human pyrin have a PP2A-B56 regulatory motif it is possible that PP2A regulates both forms through a conserved mechanism. This could be tested by inactivation of the SLiM motifs in murine and human pyrin. We have not ruled out the possible roles of other PPPs in pyrin dephosphorylation and inflammasome assembly and additional approaches will likely be needed to address this possibility. Finally, it remains to be determined if the activity of PP2A is upregulated upon RhoA inactivation, or these enzymes simply constitutively counterbalance PKN. In the case of PP2A activating NLRP3 by dephosphorylation (49), there is evidence that Bruton tyrosine kinase negatively regulates PP2A by phosphorylation of tyrosine 307 (65). It is theoretically possible that PKN could negatively regulate PPP activities by phosphorylation and additional experiments are needed to address this possibility.

## Acknowledgments

We thank Isha Nasa for help in providing advice on the phosphatase inhibitors used in Fig. S1. We would also like to thank members of the Bliska lab for providing helpful suggestions towards this project and manuscript.

## Financial support footnote

Research reported in this publication was supported by the National Institute of Allergy and Infectious Diseases of the National Institutes of Health under Award Number R01AI099222 (to JBB), and The Institute for Biomolecular Targeting (bioMT) funded by the National Institute of General Medical Sciences and grant P20-GM113132.

## References

1. Troisfontaines, P., and G. R. Cornelis. 2005. Type III secretion: more systems than you think. Physiology 20: 326–339.

2. Raymond, B., J. C. Young, M. Pallett, R. G. Endres, A. Clements, and G. Frankel. 2013. Subversion of trafficking, apoptosis, and innate immunity by type III secretion system effectors. Trends in microbiology 21: 430–441.

3. Blander, J. M., and L. E. Sander. 2012. Beyond pattern recognition: five immune checkpoints for scaling the microbial threat. Nat Rev Immunol 12: 215–225.

4. Vance, R. E., R. R. Isberg, and D. A. Portnoy. 2009. Patterns of pathogenesis: discrimination of pathogenic and nonpathogenic microbes by the innate immune system. Cell Host Microbe 6: 10–21.

5. Sollberger, G., G. E. Strittmatter, M. Garstkiewicz, J. Sand, and H. D. Beer. 2014. Caspase-1: the inflammasome and beyond. Innate immunity 20: 115–125.

6. Shin, S., and I. E. Brodsky. 2015. The inflammasome: Learning from bacterial evasion strategies. Semin Immunol.

7. Kovacs, S. B., and E. A. Miao. 2017. Gasdermins: Effectors of Pyroptosis. Trends Cell Biol.

8. Shi, J., W. Gao, and F. Shao. 2017. Pyroptosis: Gasdermin-Mediated Programmed Necrotic Cell Death. Trends Biochem Sci 42: 245–254.

9. Shi, J., Y. Zhao, K. Wang, X. Shi, Y. Wang, H. Huang, Y. Zhuang, T. Cai, F. Wang, and F. Shao. 2015. Cleavage of GSDMD by inflammatory caspases determines pyroptotic cell death. Nature 526: 660–665.

10. Bergsbaken, T., S. L. Fink, A. B. den Hartigh, W. P. Loomis, and B. T. Cookson. 2011. Coordinated host responses during pyroptosis: caspase-1-dependent lysosome exocytosis and inflammatory cytokine maturation. J Immunol 187: 2748–2754.

11. Brewer, S. M., S. W. Brubaker, and D. M. Monack. 2019. Host inflammasome defense mechanisms and bacterial pathogen evasion strategies. Curr Opin Immunol 60: 63–70.

12. Cornelis, G. R. 2006. The type III secretion injectisome. Nat Rev Microbiol 4: 811–825.

13. Viboud, G. I., and J. B. Bliska. 2005. Yersinia outer proteins: role in modulation of host cell signaling responses and pathogenesis. Annu Rev Microbiol 59: 69–89.

14. Black, D. S., and J. B. Bliska. 2000. The RhoGAP activity of the Yersinia pseudotuberculosis cytotoxin YopE is required for antiphagocytic function and virulence. Mol. Microbiol. 37: 515–527.

15. Shao, F., P. O. Vacratsis, Z. Bao, K. E. Bowers, C. A. Fierke, and J. E. Dixon. 2003. Biochemical characterization of the Yersinia YopT protease: cleavage site and recognition elements in Rho GTPases. Proc. Natl. Acad. Sci. U.S.A. 100: 904–909.

16. LaRock, C. N., and B. T. Cookson. 2012. The Yersinia virulence effector YopM binds caspase-1 to arrest inflammasome assembly and processing. Cell Host Microbe 12: 799–805.

17. Chung, L. K., Y. H. Park, Y. Zheng, I. E. Brodsky, P. Hearing, D. L. Kastner, J. J. Chae, and J. B. Bliska. 2016. The Yersinia Virulence Factor YopM Hijacks Host Kinases to Inhibit Type III Effector-Triggered Activation of the Pyrin Inflammasome. Cell Host Microbe 20: 296–306.

18. Ratner, D., M. P. Orning, M. K. Proulx, D. Wang, M. A. Gavrilin, M. D. Wewers, E. S. Alnemri, P. F. Johnson, B. Lee, J. Mecsas, N. Kayagaki, J. D. Goguen, and E. Lien. 2016. The Yersinia pestis Effector YopM Inhibits Pyrin Inflammasome Activation. PLoS Pathog 12: e1006035.

19. Brodsky, I. E., N. W. Palm, S. Sadanand, M. B. Ryndak, F. S. Sutterwala, R. A. Flavell, J. B. Bliska, and R. Medzhitov. 2010. A Yersinia Effector Protein Promotes Virulence by Preventing Inflammasome Recognition of the Type III Secretion System. Cell Host Microbe 7: 376–387.

20. Zwack, E. E., A. G. Snyder, M. A. Wynosky-Dolfi, G. Ruthel, N. H. Philip, M. M. Marketon, M. S. Francis, J. B. Bliska, and I. E. Brodsky. 2015. Inflammasome activation in response to the Yersinia type III secretion system requires hyperinjection of translocon proteins YopB and YopD. MBio 6: e02095–02014.

21. Malik, H. S., and J. B. Bliska. 2020. The pyrin inflammasome and the Yersinia effector interaction. Immunol Rev 297: 96–107.

22. Medici, N. P., M. Rashid, and J. B. Bliska. 2019. Characterization of pyrin dephosphorylation and inflammasome activation in macrophages as triggered by the Yersinia effectors YopE and YopT. Infect Immun.

23. Xu, H., J. Yang, W. Gao, L. Li, P. Li, L. Zhang, Y. N. Gong, X. Peng, J. J. Xi, S. Chen, F. Wang, and F. Shao. 2014. Innate immune sensing of bacterial modifications of Rho GTPases by the Pyrin inflammasome. Nature 513: 237–241.

24. Aubert, D. F., H. Xu, J. Yang, X. Shi, W. Gao, L. Li, F. Bisaro, S. Chen, M. A. Valvano, and F. Shao. 2016. A Burkholderia Type VI Effector Deamidates Rho GTPases to Activate the Pyrin Inflammasome and Trigger Inflammation. Cell Host Microbe 19: 664–674.

25. Schnappauf, O., J. J. Chae, D. L. Kastner, and I. Aksentijevich. 2019. The Pyrin Inflammasome in Health and Disease. Front Immunol 10: 1745.

26. Ganser-Pornillos, B. K., and O. Pornillos. 2019. Restriction of HIV-1 and other retroviruses by TRIM5. Nat Rev Microbiol 17: 546–556.

27. Weinert, C., D. Morger, A. Djekic, M. G. Grutter, and P. R. Mittl. 2015. Crystal structure of TRIM20 C-terminal coiled-coil/B30.2 fragment: implications for the recognition of higher order oligomers. Sci Rep 5: 10819.

28. Yu, J. W., T. Fernandes-Alnemri, P. Datta, J. Wu, C. Juliana, L. Solorzano, M. McCormick, Z. Zhang, and E. S. Alnemri. 2007. Pyrin activates the ASC pyroptosome in response to engagement by autoinflammatory PSTPIP1 mutants. Mol Cell 28: 214–227.

29. Chae, J. J., M. Centola, I. Aksentijevich, A. Dutra, M. Tran, G. Wood, K. Nagaraju, D. W. Kingma, P. P. Liu, and D. L. Kastner. 2000. Isolation, genomic organization, and expression analysis of the mouse and rat homologs of MEFV, the gene for familial mediterranean fever. Mamm Genome 11: 428–435.

30. Gao, W., J. Yang, W. Liu, Y. Wang, and F. Shao. 2016. Site-specific phosphorylation and microtubule dynamics control Pyrin inflammasome activation. Proc Natl Acad Sci U S A 113: E4857–4866.

31. Jones, J. D., and J. L. Dangl. 2006. The plant immune system. nature 444: 323–329.

32. Thumkeo, D., S. Watanabe, and S. Narumiya. 2013. Physiological roles of Rho and Rho effectors in mammals. Eur J Cell Biol 92: 303–315.

33. Park, Y. H., G. Wood, D. L. Kastner, and J. J. Chae. 2016. Pyrin inflammasome activation and RhoA signaling in the autoinflammatory diseases FMF and HIDS. Nat Immunol 17: 914–921.

34. Park, Y. H., E. F. Remmers, W. Lee, A. K. Ombrello, L. K. Chung, Z. Shilei, D. L. Stone, M. I. Ivanov, N. A. Loeven, K. S. Barron, P. Hoffmann, M. Nehrebecky, Y. Z. Akkaya-Ulum, E. Sag, B. Balci-Peynircioglu, I. Aksentijevich, A. Gul, C. N. Rotimi, H. Chen, J. B. Bliska, S. Ozen, D. L. Kastner, D. Shriner, and J. J. Chae. 2020. Ancient familial Mediterranean fever mutations in human pyrin and resistance to Yersinia pestis. Nat Immunol.

35. Masters, S. L., V. Lagou, I. Jeru, P. J. Baker, L. Van Eyck, D. A. Parry, D. Lawless, D. De Nardo, J. E. Garcia-Perez, L. F. Dagley, C. L. Holley, J. Dooley, F. Moghaddas, E. Pasciuto, P. Y. Jeandel, R. Sciot, D. Lyras, A. I. Webb, S. E. Nicholson, L. De Somer, E. van Nieuwenhove, J. Ruuth-Praz, B. Copin, E. Cochet, M. Medlej-Hashim, A. Megarbane, K. Schroder, S. Savic, A. Goris, S. Amselem, C. Wouters, and A. Liston. 2016. Familial autoinflammation with neutrophilic dermatosis reveals a regulatory mechanism of pyrin activation. Sci Transl Med 8: 332ra345.

36. McCoy, M. W., M. L. Marre, C. F. Lesser, and J. Mecsas. 2010. The C-terminal tail of Yersinia pseudotuberculosis YopM is critical for interacting with RSK1 and for virulence. Infect Immun 78: 2584–2598.

37. Jamilloux, Y., F. Magnotti, A. Belot, and T. Henry. 2018. The pyrin inflammasome: from sensing RhoA GTPases-inhibiting toxins to triggering autoinflammatory syndromes. Pathogens and disease 76: fty020.

38. Magnotti, F., L. Lefeuvre, S. Benezech, T. Malsot, L. Waeckel, A. Martin, S. Kerever, D. Chirita, M. Desjonqueres, A. Duquesne, M. Gerfaud-Valentin, A. Laurent, P. Seve, M. R. Popoff, T. Walzer, A. Belot, Y. Jamilloux, and T. Henry. 2019. Pyrin dephosphorylation is sufficient to trigger inflammasome activation in familial Mediterranean fever patients. EMBO Mol Med 11: e10547.

39. Simonet, M., and S. Falkow. 1992. Invasin expression in Yersinia pseudotuberculosis. Infect. Immun. 60: 4414–4417.

40. McPhee, J. B., P. Mena, and J. B. Bliska. 2010. Delineation of regions of the Yersinia YopM protein required for interaction with the RSK1 and PRK2 host kinases and their requirement for interleukin-10 production and virulence. Infect Immun 78: 3529–3539.

41. Chung, L. K., N. H. Philip, V. A. Schmidt, A. Koller, T. Strowig, R. A. Flavell, I. E. Brodsky, and J. B. Bliska. 2014. IQGAP1 is important for activation of caspase-1 in macrophages and is targeted by Yersinia pestis type III effector YopM. MBio 5: e01402–01414.

42. Loeven, N. A., A. I. Perault, P. A. Cotter, C. A. Hodges, J. D. Schwartzman, T. H. Hampton, and J. B. Bliska. 2021. The Burkholderia cenocepacia Type VI Secretion System Effector TecA Is a Virulence Factor in Mouse Models of Lung Infection. mBio 12: e0209821.

43. Chae, J. J., H. D. Komarow, J. Cheng, G. Wood, N. Raben, P. P. Liu, and D. L. Kastner. 2003. Targeted disruption of pyrin, the FMF protein, causes heightened sensitivity to endotoxin and a defect in macrophage apoptosis. Mol Cell 11: 591–604.

44. Lagrange, B., S. Benaoudia, P. Wallet, F. Magnotti, A. Provost, F. Michal, A. Martin, F. Di Lorenzo, B. F. Py, and A. Molinaro. 2018. Human caspase-4 detects tetra-acylated LPS and cytosolic Francisella and functions differently from murine caspase-11. Nature communications 9: 1–14.

45. Meerbrey, K. L., G. Hu, J. D. Kessler, K. Roarty, M. Z. Li, J. E. Fang, J. I. Herschkowitz, A. E. Burrows, A. Ciccia, and T. Sun. 2011. The pINDUCER lentiviral toolkit for inducible RNA interference in vitro and in vivo. Proceedings of the National Academy of Sciences 108: 3665–3670.

46. Case, C. L., and C. R. Roy. 2011. Asc modulates the function of NLRC4 in response to infection of macrophages by Legionella pneumophila. MBio 2: e00117–00111.

47. Pierini, R., C. Juruj, M. Perret, C. Jones, P. Mangeot, D. Weiss, and T. Henry. 2012. AIM2/ASC triggers caspase-8-dependent apoptosis in Francisella-infected caspase-1-deficient macrophages. Cell Death & Differentiation 19: 1709–1721.

48. Kofoed, E. M., and R. E. Vance. 2013. Blue native polyacrylamide gel electrophoresis to monitor inflammasome assembly and composition. Methods Mol Biol 1040: 169–183.

49. Stutz, A., C. C. Kolbe, R. Stahl, G. L. Horvath, B. S. Franklin, O. van Ray, R. Brinkschulte, M. Geyer, F. Meissner, and E. Latz. 2017. NLRP3 inflammasome assembly is regulated by phosphorylation of the pyrin domain. The Journal of experimental medicine 214: 1725–1736.

50. Nasa, I., and A. N. Kettenbach. 2018. Coordination of Protein Kinase and Phosphoprotein Phosphatase Activities in Mitosis. Front Cell Dev Biol 6: 30.

51. Aktories, K. 2011. Bacterial protein toxins that modify host regulatory GTPases. Nat Rev Microbiol 9: 487–498.

52. Namboodiripad, A. N., and M. L. Jennings. 1996. Permeability characteristics of erythrocyte membrane to okadaic acid and calyculin A. American Journal of Physiology-Cell Physiology 270: C449–C456.

53. Gavrilin, M. A., D. H. Abdelaziz, M. Mostafa, B. A. Abdulrahman, J. Grandhi, A. Akhter, A. Abu Khweek, D. F. Aubert, M. A. Valvano, M. D. Wewers, and A. O. Amer. 2012. Activation of the pyrin inflammasome by intracellular Burkholderia cenocepacia. J Immunol 188: 3469–3477.

54. Jamilloux, Y., L. Lefeuvre, F. Magnotti, A. Martin, S. Benezech, O. Allatif, M. Penel-Page, V. Hentgen, P. Seve, M. Gerfaud-Valentin, A. Duquesne, M. Desjonqueres, A. Laurent, V. Remy-Piccolo, R. Cimaz, L. Cantarini, E. Bourdonnay, T. Walzer, B. F. Py, A. Belot, and T. Henry. 2018. Familial Mediterranean fever mutations are hypermorphic mutations that specifically decrease the activation threshold of the Pyrin inflammasome. Rheumatology (Oxford) 57: 100–111.

55. Choy, M. S., M. Swingle, B. D’Arcy, K. Abney, S. F. Rusin, A. N. Kettenbach, R. Page, R. E. Honkanen, and W. Peti. 2017. PP1: Tautomycetin complex reveals a path toward the development of PP1-specific inhibitors. Journal of the American Chemical Society 139: 17703–17706.

56. Mitsuhashi, S., N. Matsuura, M. Ubukata, H. Oikawa, H. Shima, and K. Kikuchi. 2001. Tautomycetin is a novel and specific inhibitor of serine/threonine protein phosphatase type 1, PP1. Biochemical and biophysical research communications 287: 328–331.

57. Wittig, I., H. P. Braun, and H. Schagger. 2006. Blue native PAGE. Nat Protoc 1: 418–428.

58. Kreienbuhl, P., H. Keller, and V. Niggli. 1992. Protein phosphatase inhibitors okadaic acid and calyculin A alter cell shape and F-actin distribution and inhibit stimulus-dependent increases in cytoskeletal actin of human neutrophils.

59. Swingle, M., L. Ni, and R. E. Honkanen. 2007. Small-molecule inhibitors of Ser/Thr protein phosphatases. In Protein Phosphatase Protocols. Springer. 23–38.

60. Shi, Y. 2009. Serine/threonine phosphatases: mechanism through structure. Cell 139: 468–484.

61. Luheshi, N. M., J. A. Giles, G. Lopez-Castejon, and D. Brough. 2012. Sphingosine regulates the NLRP3-inflammasome and IL-1β release from macrophages. European journal of immunology 42: 716–725.

62. Jeru, I., S. Papin, S. L’Hoste, P. Duquesnoy, C. Cazeneuve, J. Camonis, and S. Amselem. 2005. Interaction of pyrin with 14.3.3 in an isoform-specific and phosphorylation-dependent manner regulates its translocation to the nucleus. Arthritis Rheum 52: 1848–1857.

63. Evdokimov, A. G., D. E. Anderson, K. M. Routzahn, and D. S. Waugh. 2001. Unusual molecular architecture of the Yersinia pestis cytotoxin YopM: a leucine-rich repeat protein with the shortest repeating unit. J Mol Biol 312: 807–821.

64. MacKintosh, C., K. A. Beattie, S. Klumpp, P. Cohen, and G. A. Codd. 1990. Cyanobacterial microcystin-LR is a potent and specific inhibitor of protein phosphatases 1 and 2A from both mammals and higher plants. FEBS letters 264: 187–192.

65. Mao, L., A. Kitani, E. Hiejima, K. Montgomery-Recht, W. Zhou, I. Fuss, A. Wiestner, and W. Strober. 2020. Bruton tyrosine kinase deficiency augments NLRP3 inflammasome activation and causes IL-1β–mediated colitis. The Journal of clinical investigation 130: 1793–1807.

