## Supplemental Figs. 1-4 for "Phosphoprotein Phosphatase Activity Positively Regulates Oligomeric Pyrin to Trigger Inflammasome Assembly in Phagocytes"

**Supplemental Figures**

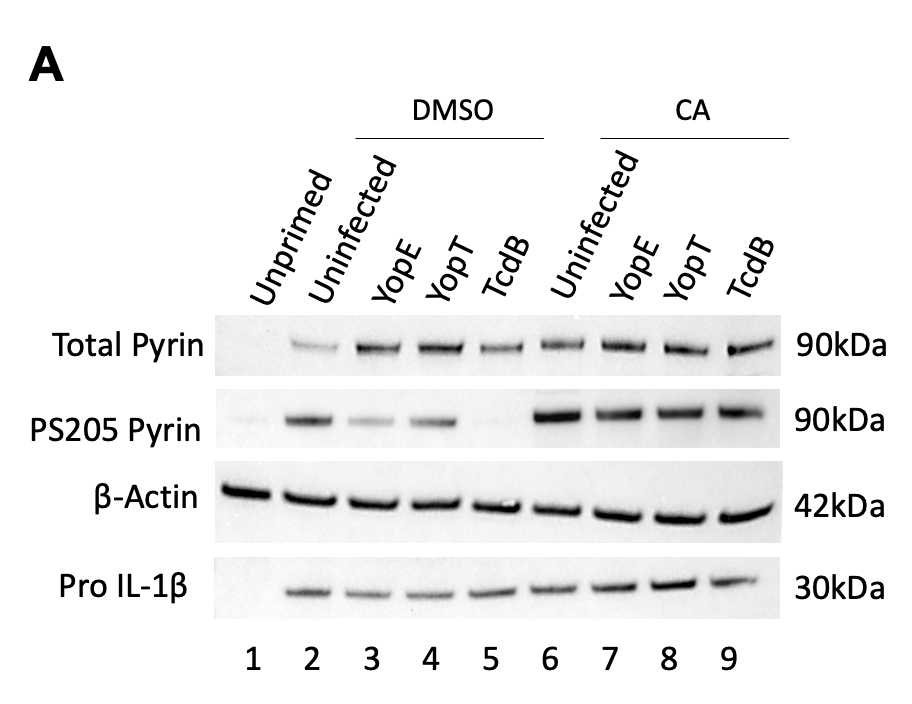

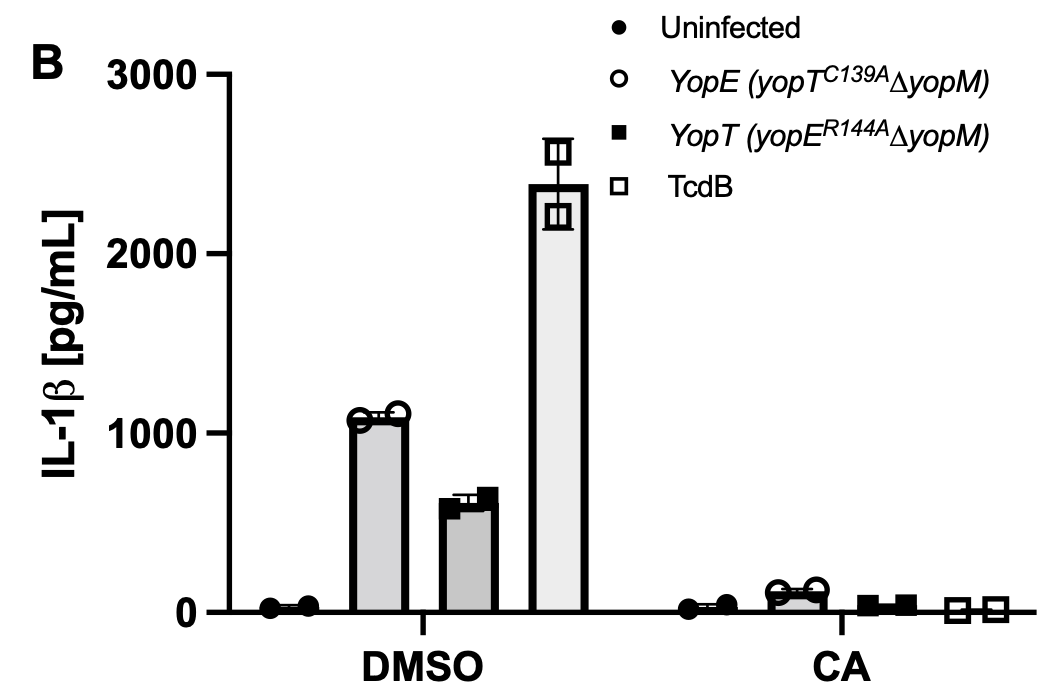

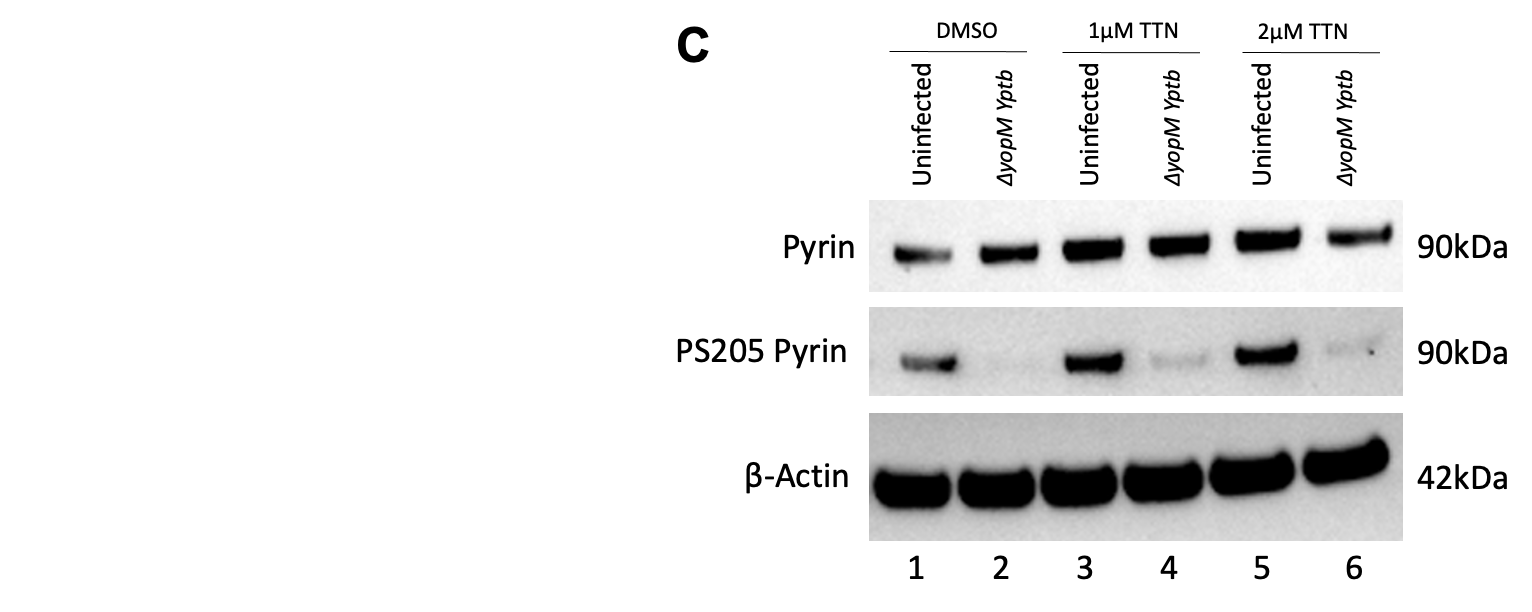

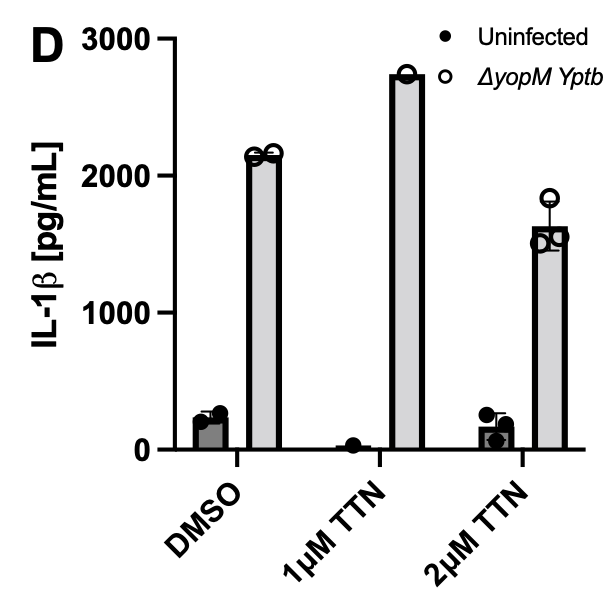

Fig.S1. PPP inhibition with CA but not TTN prevents pyrin S205 dephosphorylation and reduces IL-1β release in BMDMs infected with *∆yopM Yptb* strains. LPS-primed BMDMs were pretreated for 15 mins and maintained with either (A,B) 10nM calyculin A (CA), or (C,D) 1 or 2μM tautomycetin (TTN). In (A,B) cells were left unintoxicated or intoxicated with 0.1μg/mL of TcdB for 90 mins, or infected with either *∆yopM Yptb* YopE (*yopT^C139A^*) or YopT (*yopE^R144A^*) strains at an MOI of 30 for 90min. In (C,D) cells were left uninfected or infected with *∆yopM Yptb* at an MOI of 30 for 90 mins. Unprimed BMDMs or treatment with vehicle DMSO were used as controls. A,C) Immunoblot analysis of total and PS205 pyrin or pro-IL-1β in BMDM lysates. Actin was used as a loading control. B,D) Mature IL-1β in supernatants as quantified by ELISA. In (B) each data group is presented as an average (error bars are standard deviation) of two technical replicates from one experiment. In (D) each data group is presented as an average (error bars are standard deviation) of technical replicates from one experiment.

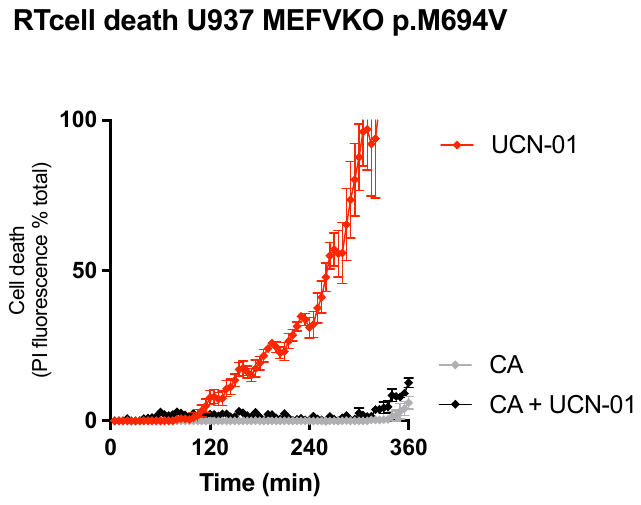

Fig.S2. CA inhibits UCN-01-mediated cell death in human monocytes expressing M694V pyrin variant. U937 *MEFV*^KO^ cells expressing p.M694V *MEFV* variant were treated with UCN-01 (12.5 µM) with or without pre-treatment with Calyculin A (40 nM). Propidium iodide (PI) influx/fluorescence was monitored every 5 min for 6h. Each data group is presented as an average (error bars are standard deviation) of three technical replicates.

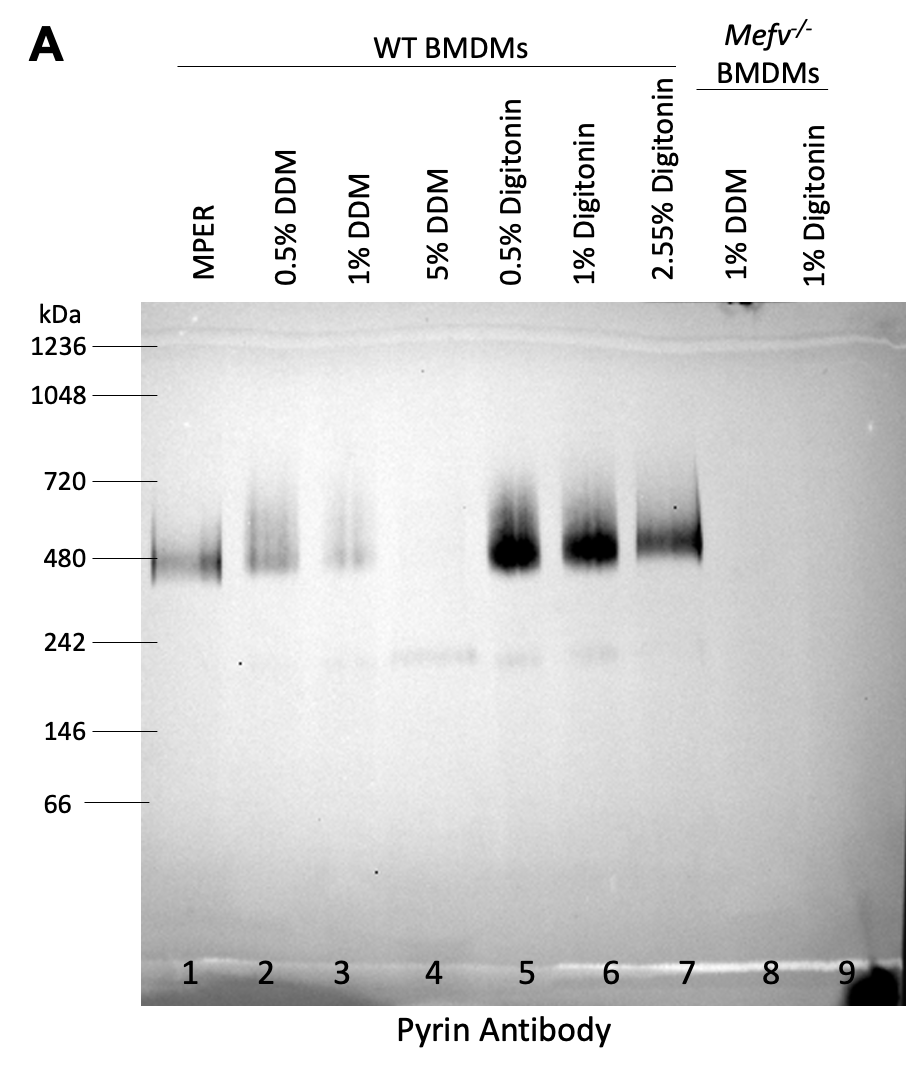

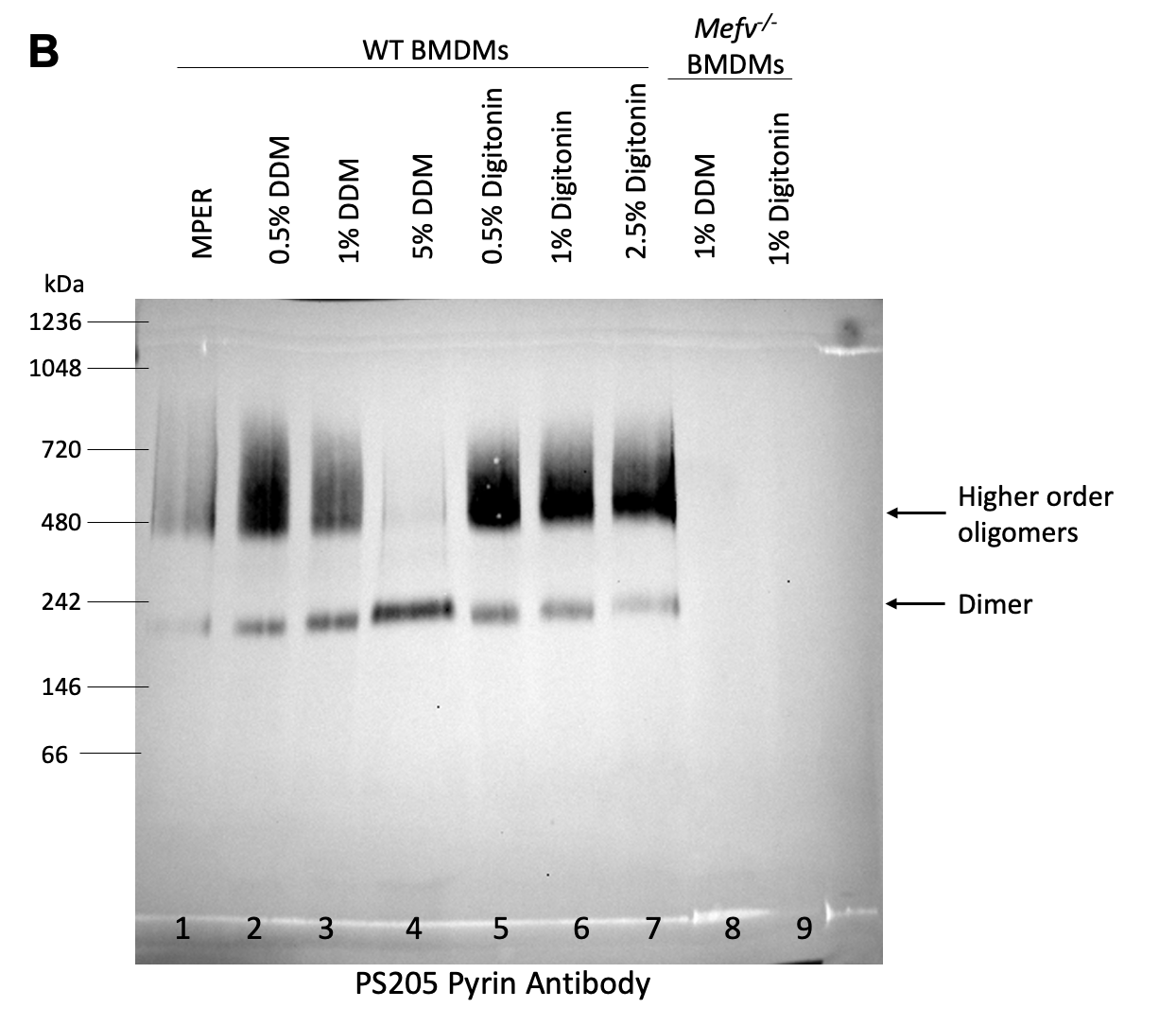

Fig.S3. BMDM lysates prepared with digitonin are optimal for detection of pyrin oligomers using BN-PAGE and immunoblotting. LPS-primed WT and *Mefv*^-/-^ BMDMs were lysed in various non-ionic detergent concentrations as indicated. Lysate samples were separated by BN-PAGE and immunoblotted to visualize total (A) and PS205 pyrin (B) oligomers. Positions of BN-PAGE molecular weight standards are shown on the left of each panel, and dimer and higher order oligomer are indicated on the right of (B).

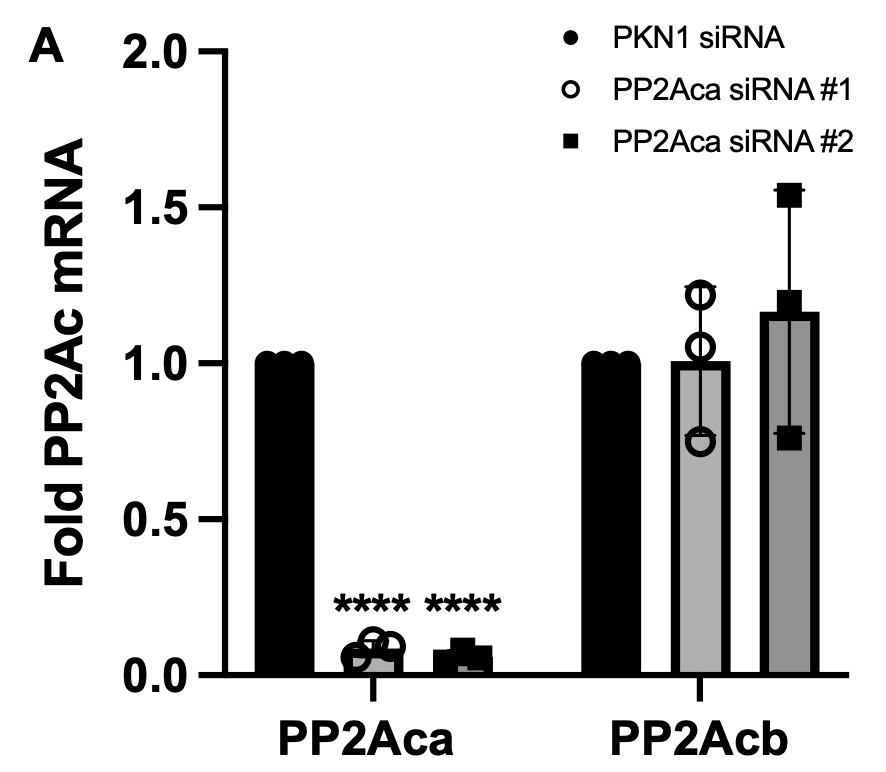

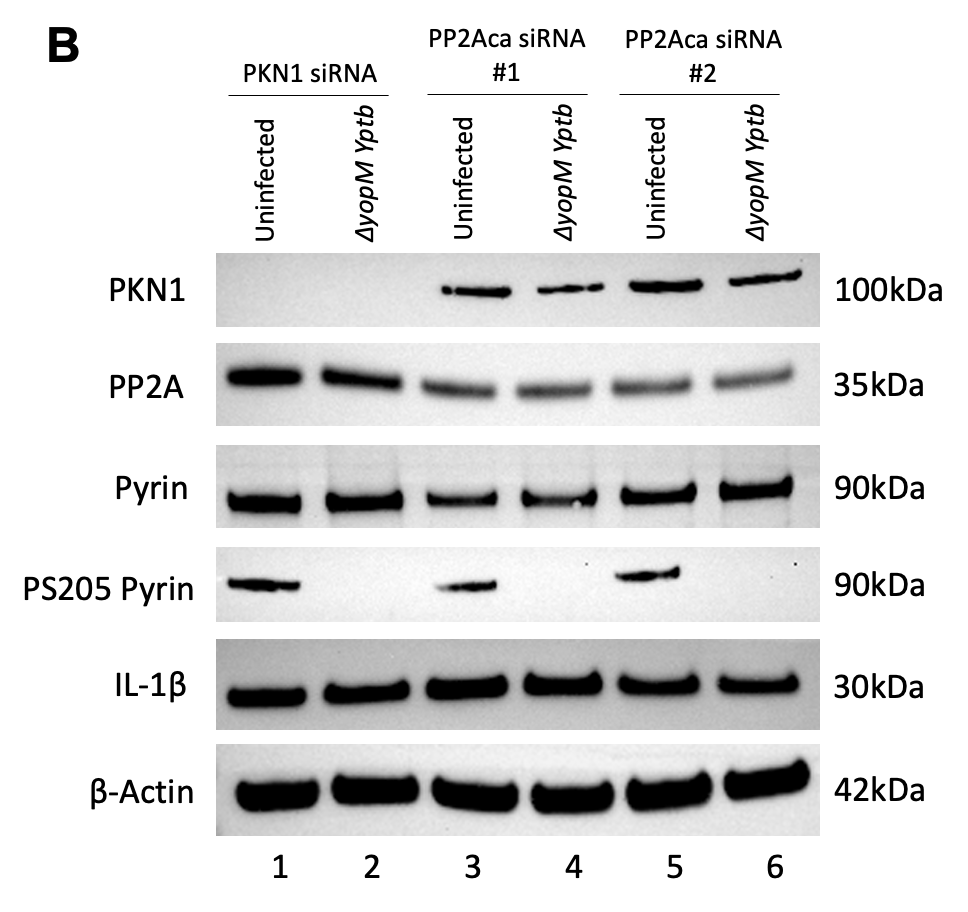

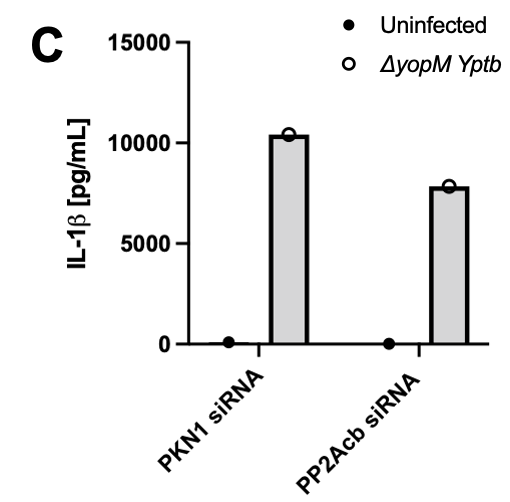

Fig.S4. Individual siRNA knock-down of PP2Aca and PP2Acb in BMDMs does not prevent pyrin inflammasome activation. WT BMDMs were electroporated with siRNAs targeting PKN1 (400pmol), PP2Aca (1000pmol), or PP2Acb (500pmol). 24hr after electroporation the BMDMs were LPS-primed. 48hr following electroporation, cells were either left uninfected or infected with *∆yopM Yptb* at an MOI of 30 for 90 mins. A) RT-qPCR analysis of mRNA transcripts of PP2Aca and PP2Acb in uninfected BMDMs. Results were normalized to Hprt mRNA levels. Each data group is presented as an average (error bars are standard deviation) of three independent experiments. One-way ANOVA was applied to calculate significance and p-values as compared to PKN1 control siRNA are indicated. P-value<0.05 was considered significant; <0.0001 (****). B) Immunoblot analysis of PKN1, PP2A, total and PS205 pyrin, and pro-IL-1β in uninfected and *∆yopM Yptb-*infected BMDM lysates. Actin was used as a loading control. C) Mature IL-1β in supernatants as quantified by ELISA from one experiment.
